# A regulatory T cell signature provides a shared molecular basis for the therapeutic window of opportunity in rheumatic disease

**DOI:** 10.64898/2026.09.21.753213

**Authors:** Marie Binvignat, Federica Martina, Fabien Pitoiset, Alexandra Roux, Roberta Lorenzon, Pierre Hodara, Pierre Barennes, Hélène Vantomme, Michèle Sastre, Signe Hassler, Claire Ribet, Alice Courties, Vanessa Mhanna, Paul Stys, Leslie Adda, Vimala Diderot, Eric Vicaut, Caroline Aheng, Nicolas Coatnoan, Adrien Six, Nicolas Tchitchek, Michelle Rosenzwajg, Francis Berenbaum, David Klatzmann, Jérémie Sellam, Encarnita Mariotti-Ferrandiz

## Abstract

Rheumatic diseases, including rheumatoid arthritis (RA), spondyloarthritis (SpA) and osteoarthritis (OA), show distinct phenotypes yet respond to overlapping therapies, implicating shared immune mechanisms. In the Trans*i*mmunom cohort, we profiled peripheral blood from 240 individuals (47 healthy, 44 OA, 91 RA, 58 SpA) across deep immunophenotyping, immunoproteomics and Treg-Teff transcriptomics. Single-layer analyses revealed broader Treg than Teff remodeling, along with a shared pattern of reduced activated Tregs and expanded Helios⁺ Tregs across all diseases, alongside a decrease in functional Treg subpopulations, including CTLA4⁺ and CD45RA⁻ Tregs. In RA specifically, LAG3⁺ Tregs were also expanded. Combining omics layers outperformed single-layer approaches for disease classification. Among individual layers, Treg transcriptomes were most discriminative, and integration uncovered disease-specific programs. Unsupervised clustering identified a cross-disease cluster independent of activity, treatment and age, mapping to early disease (≤2 years) and dominated by a Treg dysfunction-associated program. These results provide a biological rationale for the therapeutic “window of opportunity” concept and duration-stratified Treg-directed trials.

## INTRODUCTION

Rheumatic diseases differ in clinical presentation, radiographic features, and treatment response, yet share a defining paradox: distinct diagnoses respond to the same targeted therapies^1,2^. Conventional, biologic and targeted-synthetic disease-modifying antirheumatic drugs (DMARDs) are effective across diseases long considered mechanistically unrelated, implying that overlapping inflammatory pathways drive clinically divergent phenotypes^3–6^. Yet predicting treatment response and disease trajectory remains the central challenge in rheumatology. The therapeutic “window of opportunity” is well established: early intervention yields deeper, more durable responses, including drug-free remission, whereas delayed response often leads to self-sustaining, refractory inflammation^7–10^. Among these overlapping pathways, the balance between regulatory T cells (Tregs) and effector T cells (Teffs) is central to immune homeostasis and a plausible common denominator across diseases. Treg (CD4+FOXP3+, CD25-high and CD127-low) are indispensable for self-tolerance^11^. They restrain inflammation through IL-2 consumption, CTLA-4–mediated control of antigen-presenting cells, inhibitory cytokines (IL-10, TGF-β, IL-35), metabolic disruption and cytolysis, and they contribute to tissue repair^12^. When Tregs fail in number or function, or when Teffs resist suppression, the equilibrium tips toward chronic inflammation and autoimmunity^13^. This axis is disturbed across rheumatic diseases, but in distinct ways; in rheumatoid arthritis (RA), Treg suppressive function tracks inversely with disease activity and is restored in clinical remission^14^. In spondyloarthritis (SpA), Treg biology is tied to the IL-23/IL-17 axis, with plasticity toward a Th17-like state at sites of enthesitis and gut inflammation^15^, and more recently, in osteoarthritis (OA), Treg dysfunction profile have been linked to clinical features including knee pain^16^

Rheumatic diseases arise from interactions that span molecular scales (cellular, proteomic, and transcriptional) and no single measurement captures them thoroughly. Each omics layer offers a partial view, and disease-relevant biology may reside in the relationships between them. Integrative multi-omics can expose these inter-omic interactions, separate shared from disease-specific signatures, and generate testable hypotheses.

Here, within the Trans*i*mmunom cohort, we present an integrated Treg–Teff atlas spanning 240 individuals with RA, SpA, OA and healthy volunteers (HV). By combining deep immunophenotyping, immunoproteomics (cytokines and chemokines), and transcriptomics of sorted Tregs and Teffs with integrative multi-omics methods (multi-block sparse partial least-squares discriminant analysis (sPLS-DA), we aimed to identify molecular patterns that stratify patients and delineate the shared and disease-specific immunology of rheumatic disease.

## RESULTS

### Identification of a Treg-Teff atlas across rheumatic diseases

To build an immunological Treg–Teff atlas of rheumatic disease, we profiled peripheral blood from 240 participants in the Trans*i*mmunom study (NCT02466217): 47 HV, 44 patients with knee OA, 91 with RA and 58 with SpA (**Fig. 1a**). Clinical characteristics of the cohort are summarized in **Table 1**; inclusion criteria and detailed clinical information of RA and SpA patients are displayed in **Tables S1–S3**. Each sample was profiled across four complementary layers. Deep immunophenotyping, using nine flow cytometry panels developed in-house, resolved *137* cell populations, including T cells, B cells, NK cells and monocytes subpopulations (**Fig. S1, Table S4**). Immunoproteomics quantified 62 cytokines and chemokines by Luminex (**Table S5**). Transcriptomic profiling was performed by bulk mRNA sequencing of two sorted CD4⁺ T cell compartments: regulatory T cells (Treg; CD4⁺CD25⁺CD127ˡᵒ) and effector T cells (Teff; CD4⁺CD25⁻CD127⁺). To enable cross-disease integration, we first performed compartment-specific differential expression analyses by comparing each disease with HV.

**Fig. 1.**
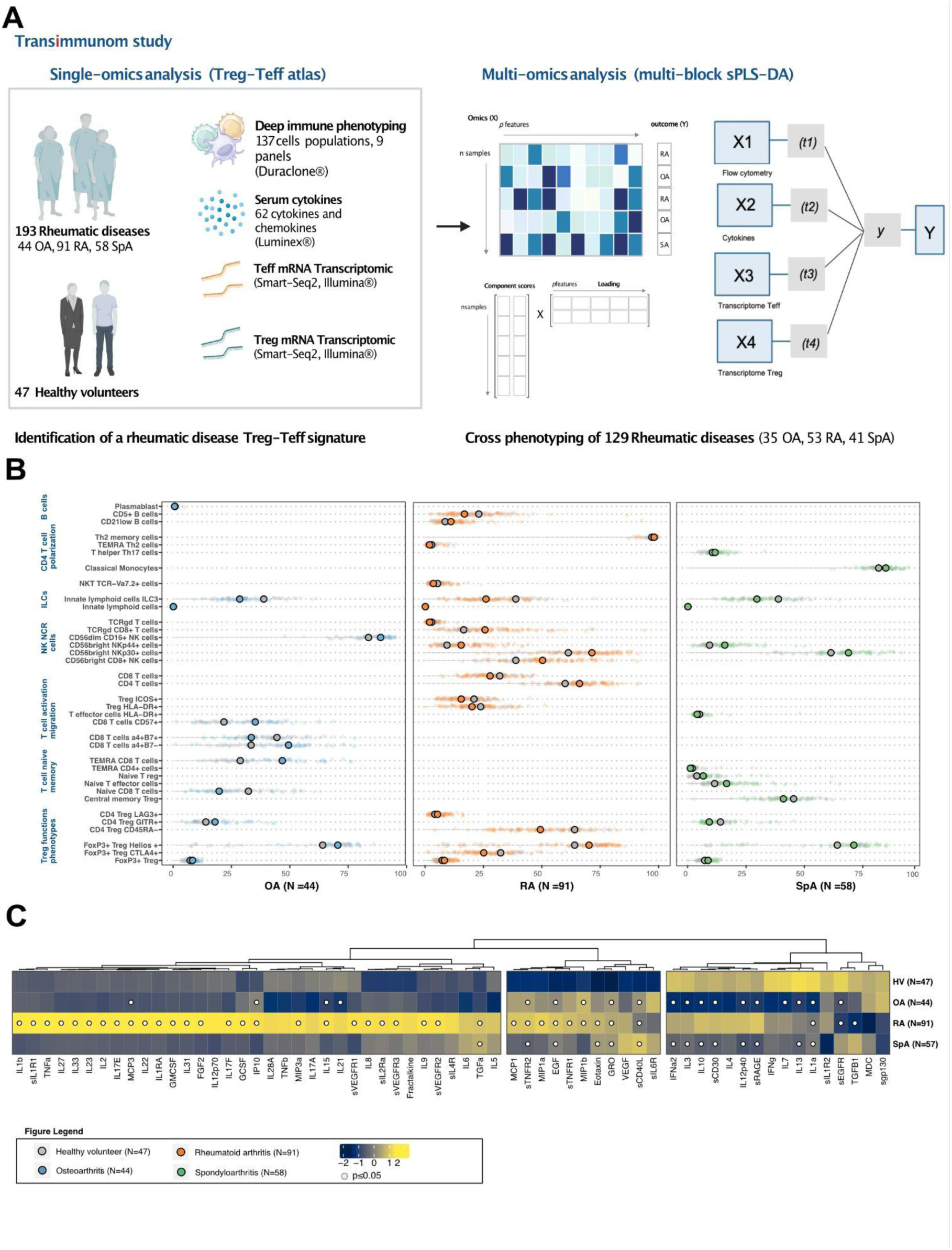
Overview of the Trans*i*mmunom study and identification of a Treg–Teff atlas across rheumatic diseases. **A.** Study design. Peripheral blood from 240 participants of the Trans*i*mmunom cohort 47 healthy volunteers (HV), 44 osteoarthritis (OA), 91 rheumatoid arthritis (RA) and 58 spondyloarthritis (SpA) was profiled across four complementary layers: deep immunophenotyping by flow cytometry (9 panels, 137 cell populations), immunoproteomics (62 serum cytokines and chemokines by Luminex®), and transcriptomics of two sorted CD4⁺ T cell compartments, regulatory (Treg; CD4⁺CD25⁺CD127ˡᵒ) and effector (Teff; CD4⁺CD25⁻CD127hi) T cells. Data were analysed both within each layer and by integrative multi-omics. **B.** Flow cytometry. Cell populations differing significantly from HV in at least one disease (Mann–Whitney U test, Benjamini–Hochberg FDR ≤ 0.05) are shown. For each population, the median proportion is shown per group (HV, OA, RA, SpA); only populations significantly different from HV are represented (FDR ≤ 0.05). **C.** Immunoproteomics. Heatmap of median serum cytokine and chemokine abundance per group. Cytokines reaching significance compared to HV in a given disease are marked with a dot (FDR ≤ 0.05).

**Table 1.** Clinical characteristic of Trans*i*mmunom patients (N=240). Patients are stratified by diseases including healthy volunteers (N=47), osteoarthritis (N=44), rheumatoid arthritis (N=91), spondyloarthritis (N=58). Continuous variables were expressed as mean and standard deviation (SD) in each subgroup, while categorical variables were expressed as percentages. One-way ANOVA and chi-square tests were applied to assess the differences between diseases. A significant p-value was defined with a threshold of ≤ 0.05. Missing values are expressed as a percentage (%). For disease duration and early-disease proportion, statistics and missing values were computed only for the rheumatic disease groups.

|  | Healthy volunteer<br>(N=47) | Osteoarthritis<br>(N=44) | Rheumatoid Arthritis<br>(N=91) | Spondyloarthritis<br>(N=58) | p-value | missing<br>(%) |
| --- | --- | --- | --- | --- | --- | --- |
| **Age** (years)<br>(mean (SD)) | 38.79 (12.88) | 65.23 (9.88) | 49.71 (13.44) | 39.50 (12.08) | <0.001 | 0 |
| **BMI** (kg/m<sup>2</sup>)<br>(mean (SD)) | 24.19 (3.38) | 29.17 (7.15) | 25.76 (5.34) | 25.41 (4.57) | <0.001 | 2.9 |
| **Sex** (%) Female | 24 (51.1) | 29 (65.9) | 74 (81.3) | 27 (46.6) | <0.001 | 0 |
| Male | 23 (48.9) | 15 (34.1) | 17 (18.7) | 31 (53.4) |  |  |
| **Lymphocytes**<br>(/mm<sup>3</sup>)<br>(mean (SD)) | 1.67 (0.44) | 1.94 (0.53) | 2.39 (3.34) | 2.18 (0.71) | ns | 1.7 |
| **hs-CRP** (mg/L)<br>(mean (SD)) | 1.15 (1.01) | 2.77 (2.83) | 10.48 (26.06) | 11.82 (19.10) | 0.005 | 12.1 |
| **Disease duration**<br>(years)<br>(mean (SD)) | NA | 8.44 (8.98) | 5.30 (7.52) | 4.81 (7.37) | 0.047 | 0.5 |
| **Early diseases**<br>(Yes (%)) | NA | 9 (20.9) | 38 (41.8) | 32 (55.2) | 0.04 | 0.5 |

We first compared the proportion of each cell population between HV and each disease group (Mann–Whitney U test with Benjamini–Hochberg correction). Seventy-four populations differed at nominal significance (P < 0.05) and 36 after correction (FDR ≤ 0.05) in at least one disease (**Fig. 1b**, **Table S6**). Most cell populations were T related, with few B and NK cell subsets associated with RA. In RA (n = 91), three functional Treg subsets were reduced relative to HV (n = 47; values are median [IQR]): CD45RA⁻ CD4⁺ Tregs, 49.39% [37.00–62.24] versus 64.00% [51.38–70.71] (FDR = 0.0015); LAG3⁺ CD4⁺ Tregs, 5.34% [3.94–7.80] vs 4.22% [4.06–4.58] versus (FDR = 0.0091); and CTLA4⁺ CD4⁺ Tregs, 25.14% [20.65–30.52] versus 32.40% [25.75–40.22] (FDR = 0.0015). The Helios⁺ compartment moved in the opposite direction: Helios⁺ FoxP3⁺ Tregs were expanded across all three diseases relative to HV (64.07% [23.54–72.43]), reaching 70.55% [66.63–74.95] in OA (FDR = 0.0171), 70.24% [59.20–77.19] in RA (FDR = 0.0065) and 71.15% [65.06–76.39] in SpA (FDR = 0.0103). Taken together, these results reveal a shared pattern of Treg alteration across rheumatic diseases, including decrease of activated Tregs and increase in LAG3+ and Helios+ Tregs.

Serum cytokine profiling in 239 of the 240 participants (HV, n = 47; OA, n = 44; RA, n = 91; SpA, n = 57; one SpA sample was unavailable for cytokine profiling) identified 51 cytokines at P < 0.05 and 47 at FDR ≤ 0.05 that distinguished at least one disease from HV (Mann–Whitney U test, Benjamini–Hochberg correction; **Fig. 1c, Table S7**). RA carried the strongest signature, with 38 differentially abundant cytokines, compared with 19 in OA and 14 in SpA.

The RA profile combined a proinflammatory shift with disruption of Treg-associated cytokines, including Tumor Necrosis Factor (TNF) increase in RA (72.59 MFI [58.26–88.83]) relative to HV (62.79 [52.57–73.60]; FDR = 0.049), consistent with the inflammatory hallmark of the disease. In parallel, the two cytokines most closely tied to Treg cell biology moved in opposite directions: IL-2 increased in RA (23.35 [7.74–90.14] versus 9.01 [5.78–25.49]; FDR = 0.004), whereas TGF-β declined (1,918.36 [1,869.92–2,058.76] versus 2,073.08 [2,019.13–2,105.12]; FDR < 0.001). This pairing (elevated IL-2 alongside reduced TGF-β) points to a perturbed rather than simply diminished Treg cytokine environment, complementing the contraction of functional Treg subsets seen by flow cytometry. Consistent with alterations in the Treg compartment, circulating IL-10 levels were reduced in OA and SpA. Compared with HV (22.81 [16.06–28.27]), IL-10 decreased to 10.28 [6.83–22.69] in OA (FDR < 0.001) and 15.20 [8.36–29.50] in SpA (FDR = 0.035).

We next performed differential gene expression analysis separately in Teff and Treg cells, comparing patients with RA (Teff n = 82, Treg n = 59), OA (Teff n = 44, Treg n = 35) and SpA (Teff n = 51, Treg n = 45) against HV (Teff n = 38, Treg n = 22). Genes were called differentially expressed at FDR ≤ 0.05, |log₂(fold change)| ≥ log₂(2) and baseMean ≥ 8 (**Fig. 2a** and **Supplementary Figs. 3–9, Table S8**). In Teff cells, upregulated genes were largely disease specific. Relative to HV, RA showed 51 upregulated genes (including *IFNG*, *CCL3* and *IFI27*) and 49 downregulated genes; SpA showed 53 upregulated genes (including *S100A8*, *S100A9*, *IL31RA* and *CXCR2*) and 26 downregulated genes; and OA showed 54 upregulated genes (such as *IL6*, *KLRD1*, *GZMB* and *ITGB3*) and 26 downregulated genes. Whereas the upregulated programs were largely distinct between diseases, the downregulated program was shared across conditions and included *CCR5*, *IL3RA*, *SMAD7* and *CCR2*. In Treg cells, transcriptional changes were dominated by shared downregulation. Relative to HV, RA showed 26 upregulated and 48 downregulated genes (including *HLA-DRB3*, *HLA-DQA2* and *FOXB1*); OA showed 9 upregulated and 176 downregulated genes (the latter including *CCL4, IRF8, CXCL2, SOCS6, CCL20, TGFBI and TNFRSF12A)*; and SpA showed 41 upregulated genes (including *TNFSF4, ITGB5* and *CD160*) and 35 downregulated genes.

**Fig. 2.**
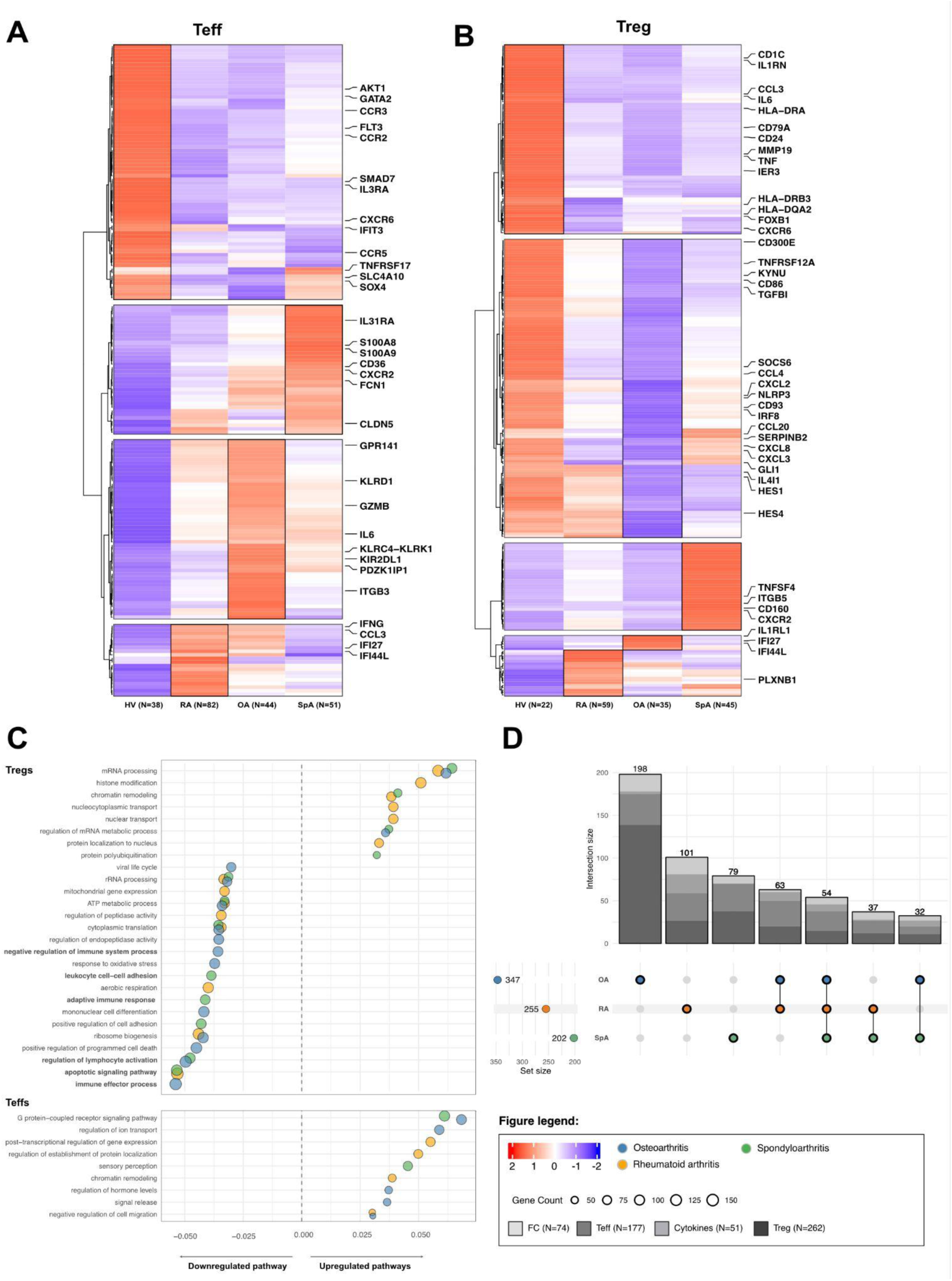
Differential expression and functional enrichment in Treg and Teff cells across rheumatic diseases. **A-B.** Differential gene expression. Mean expression (normalized counts) per disease group for genes differentially expressed relative to HV, shown separately for (A) effector (Teff) and (B) regulatory (Treg) T cells. Genes were called differentially expressed at FDR ≤ 0.05, |log₂ fold change| ≥ 1 and baseMean ≥ 8 (Teff: HV n = 38, OA n = 44, RA n = 82, SpA n = 51; Treg: HV n = 22, OA n = 35, RA n = 59, SpA n = 45). **C.** Over-representation analysis. Enriched biological pathways among the up- and down-regulated differentially expressed genes in each disease, shown separately for Treg and Teff cells (pathways retained at FDR ≤ 0.05, gene ratio ≥ 0.03 and count ≥ 20). **D.** UpSet plot summarizing the features unique to, or shared across, OA, SpA and RA, aggregated over all omics layers (flow cytometry, cytokines, Treg and Teff transcriptomes).

In Over-Representation Analysis (ORA) of the differentially expressed genes (p ≤ 0.05, |log₂FC| > 0, q ≤ 0.02), we retained pathways at FDR ≤ 0.05, gene ratio ≥ 0.03 and count ≥ 20) (**Fig. 2c** and **Table S10**). In Teff cells, enriched pathways were predominantly upregulated: 4 in RA, 2 in SpA and 5 in OA, the latter including G-protein-coupled receptor signaling pathway and negative regulation of cell migration. In Treg cells, the pattern was dominated by pathway downregulation across diseases. RA showed 6 upregulated and 8 downregulated pathways, the downregulated set including rRNA processing, ATP metabolic process and apoptotic signaling. SpA showed 4 upregulated and 8 downregulated pathways, with downregulation of regulation of lymphocyte activation, positive regulation of cell adhesion and adaptive immune response. OA showed 2 upregulated pathways, including the regulation of mRNA metabolism, and 12 downregulated pathways, including negative regulation of immune system process and immune effector process.

Across all layers, we identified 564 differential features: 74 cell populations by flow cytometry and 51 cytokines (both P < 0.05), together with 262 genes in Treg and 177 in Teff (**Fig. 2d, Fig S10**). Of these, 198 were specific to OA, 101 to RA and 79 to SpA, 186 were shared by at least two diseases, including 54 common to all three. Collectively, our results indicate that Tregs undergo broader remodeling than Teffs across rheumatic diseases, with a predominant loss of pathways involved in immune regulation, activation and cellular homeostasis.

### Multi-omics outperform single omics in disease classification and Treg alteration drive separation of rheumatic diseases

Having mapped the shared and disease-specific molecular and cellular signatures within each omics layer, we next sought to integrate these modalities into a unified multi-omics model to define cross-disease molecular patterns. To this end, we built a multi-block sparse Partial Least Square Discriminant Analysis (sPLS-DA) model on 129 patients for which complete data across all four layers were available (OA n = 35, SpA n = 41, RA n = 53). The model retained 209 features overall (68 cell subsets, 16 cytokines, 100 transcripts in Tregs and 25 in Teffs), and we assessed discrimination against models trained on each layer alone. Model hyperparameters, fine tuning and classification are described in the Methods. Performance was estimated by five-fold cross-validation repeated 10 times, using the averaged consensus prediction across layers.

The integrated model gave the best overall separation of diseases. Projection of individuals onto the first two components separated the groups more clearly for the averaged consensus than for any single block (**Fig. 3a**), and the cross-validated overall error rate was 0.30 (SD 0.03), lower than any single block (cell subsets 0.45 (0.04), cytokines 0.39 (0.03), Treg transcripts 0.41 (0.03) and Teff transcripts 0.44 (0.04)) (**Table 2**, **Fig. S11-12**). The outperformance of integration was clearest for RA and SpA, classified at error rates of 0.20 (0.04) and 0.35 (0.06), respectively. The weighted consensus prediction was used during hyperparameter tuning, as it accounts for the correlation between blocks. However, because this weighting is defined only once blocks are combined, it does not permit decomposition into per-block error rates. Under this scheme the overall error rate was higher, at 0.42 (0.04), but the overall and balanced error rates were near-identical (both 0.42 [0.04]), arguing against a strong effect of class imbalance on the final prediction (**Table S12**). Per-block one-versus-rest AUCs were consistent with the error rates. The Treg transcripts was the strongest single block, distinguishing each disease from the others with high accuracy (OA vs rest AUC 0.855; SpA vs rest 0.948; RA vs rest 0.916). Flow cytometry discriminated OA most effectively (AUC 0.916), whereas cytokines were informative only for RA and SpA (AUC 0.800 and 0.770) and near-random for OA (AUC 0.560, P = 0.27) (**Table S11, Fig. S13**).

**Fig. 3.**
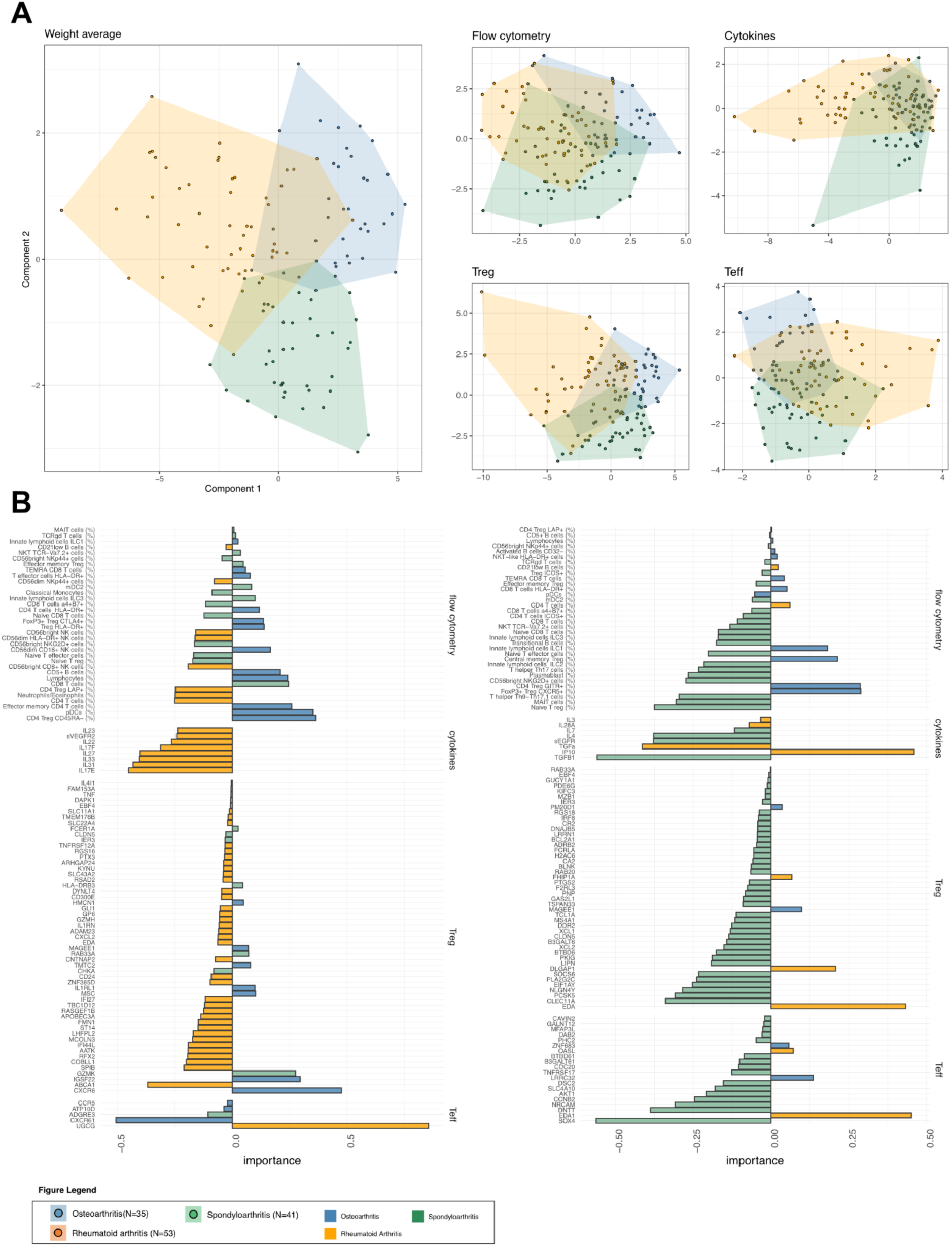
Single-omics versus integrative multi-omics discrimination of patient groups. **A.** Individual sample plots on the first two components, component 1 (horizontal axis) and component 2 (vertical axis), for the integrative multi-omics model (left panel) and for each of the four single-omics blocks, showing the distribution of patients in the reduced component space. **B.** Corresponding loading plots indicating the contribution of individual features to each component.

**Table 2.**
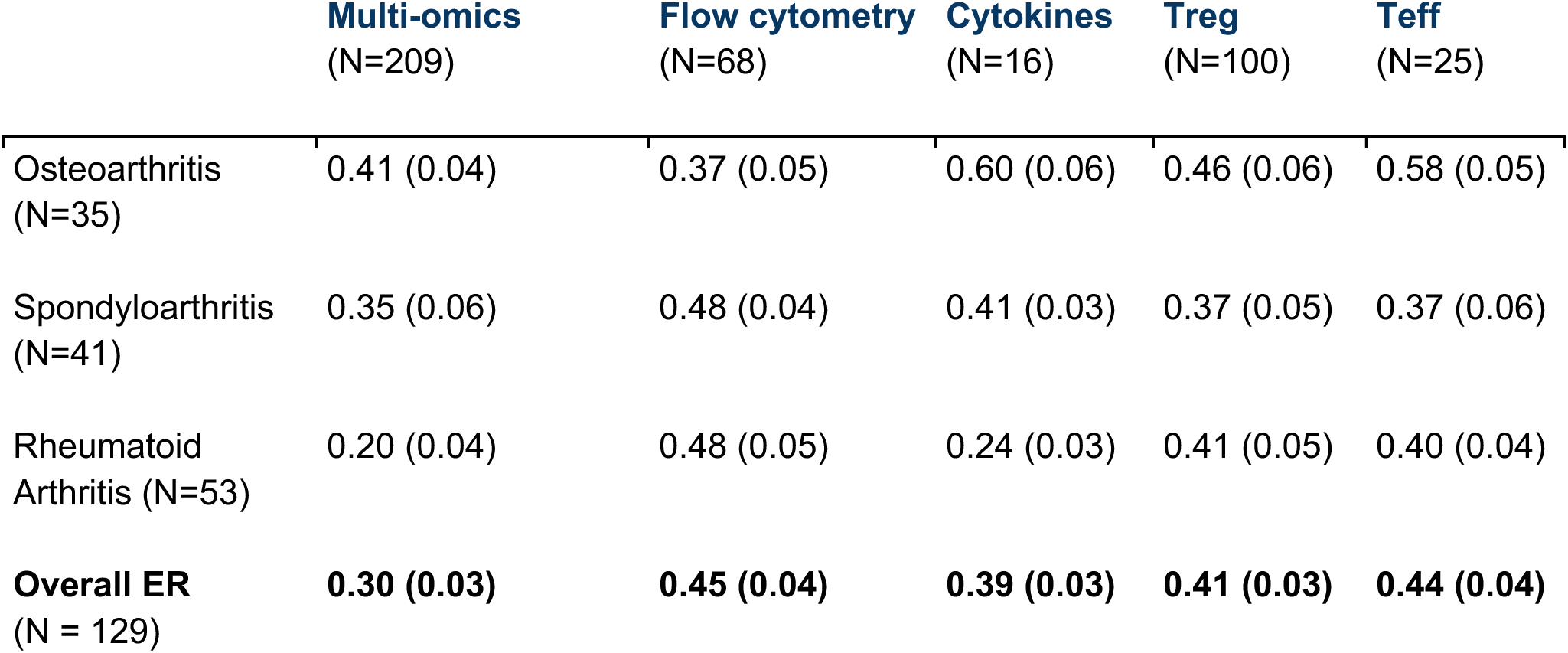
Classification error rates of multi-omics integration versus single-block models across rheumatic diseases. Classification performance of the integrative multi-omics model (N=209 features) compared with individual data blocks: flow cytometry (N=68), cytokines (N=16), Tregs (N=100), Teffs (N=25). Error rates (ER) are presented as mean (SD) for each disease class, osteoarthritis (N=35), spondyloarthritis (N=41), rheumatoid arthritis (N=53) and overall (N=127).

|  | **Multi-omics**<br>**(N=209)** | **Flow cytometry**<br>**(N=68)** | **Cytokines**<br>**(N=16)** | **Treg**<br>**(N=100)** | **Teff**<br>**(N=25)** |
| --- | --- | --- | --- | --- | --- |
| Osteoarthritis<br>(N=35) | 0.41 (0.04) | 0.37 (0.05) | 0.60 (0.06) | 0.46 (0.06) | 0.58 (0.05) |
| Spondyloarthritis<br>(N=41) | 0.35 (0.06) | 0.48 (0.04) | 0.41 (0.03) | 0.37 (0.05) | 0.37 (0.06) |
| Rheumatoid<br>Arthritis (N=53) | 0.20 (0.04) | 0.48 (0.05) | 0.24 (0.03) | 0.41 (0.05) | 0.40 (0.04) |
| **Overall ER**<br>**(N = 129)** | **0.30 (0.03)** | **0.45 (0.04)** | **0.39 (0.03)** | **0.41 (0.03)** | **0.44 (0.04)** |

Component loadings identified the features driving separation along the first two dimensions (**Fig. 3b**, **Tables S12–S13**). Integrative multi-omics modelling retained 102 features on Component 1 (34 cell subsets, 8 cytokines, 55 Treg and 5 Teff transcripts) and 107 features on Component 2 (34 cell subsets, 8 cytokines, 45 Treg and 20 Teff transcripts), and 27 features were common in both components.

Within each component, features were assigned to the disease group in which their mean abundance or expression was highest, and the two components captured largely distinct disease axes. Component 1 was dominated by RA-associated features (57 of 102), which spanned every modality but were most concentrated in the Treg transcriptome (40 of 55 transcripts) and the cytokine panel (8 of 8 cytokines). This RA cytokine signature was internally coherent, comprising the Th17/IL-1-family axis (IL-17F, IL-22 and IL-23) together with the epithelial alarmins IL-25 (IL17E), IL-31 and IL-33, as well as IL-27 and sVEGFR2, and was accompanied in the Treg compartment by a type-I-interferon-stimulated gene module (IFI44L, IFI27 and RSAD2). The remaining features were split between OA (n = 22) and SpA (n = 23). OA-assigned features were enriched for regulatory and effector-memory CD4⁺ subsets (CD4 Treg CD45RA⁻, effector-memory CD4 T cells and FoxP3⁺ Treg CTLA4⁺) and for CXCR6, the top-ranked OA feature in both the Treg and Teff compartments, whereas naive and NK-cell subsets contributed to SpA discrimination. Component 2 was strongly SpA-driven (80 of 107 features), with SpA accounting for the majority of Treg (40 of 45) and Teff (16 of 20) transcripts and for 20 of 34 cell subsets. The SpA cellular profile was distinguished by mucosal and type-3 immunity, including MAIT cells, Th17 and Th9/Th17.1 helper cells and ILC2/ILC3 populations. At the cytokine level, TGF-β1 was the leading contributor in SpA, whereas IP-10 (CXCL10) and TGF-α were highest in RA. Notably, the follicular-like regulatory subset FoxP3⁺ Treg CXCR5⁺, among the highest-ranked cell features on this component, was markedly lower in SpA than in OA or RA, indicating relative depletion of this population in spondyloarthritis.

Together, these results show that integrating cellular, proteomic and transcriptomic data yields a more accurate patient stratification than single omic analysis, and identifies regulatory T cell programs as the most discriminating dimension.

### Identification of a multi-omics immunological signature associated with RA, SA and OA

Across the 182 multiomics features that discriminated patients from HV, we grouped features and patients by hierarchical clustering (Ward’s D2 linkage) on a correlation-based distance (1 − Pearson correlation), resolving an immunological footprint for each patient. Hierarchical clustering of patients identified four distinct clusters (N = 29, 24, 21 and 55; **Fig. 4** and **Fig. S14**). Three of these clusters mapped closely onto a single diagnosis: cluster 2 was enriched for RA (N=23/24, 95.8%) and defined by a strong cytokine signature, cluster 3 for OA (N=18/21, 85.7%) and cluster 4 for SpA (29/55, 52.7%). In contrast, cluster 1 comprised a mixture of RA (14/29 48.3%), SpA (N=11/29 37.9%) and OA (N=4/29 13.8%) patients and was instead driven by a pronounced transcriptomic Treg signature. Cluster compositions are detailed in **Fig. 5a-b**, **Fig S15-17**, **Table S14-15.**

**Fig. 4.**
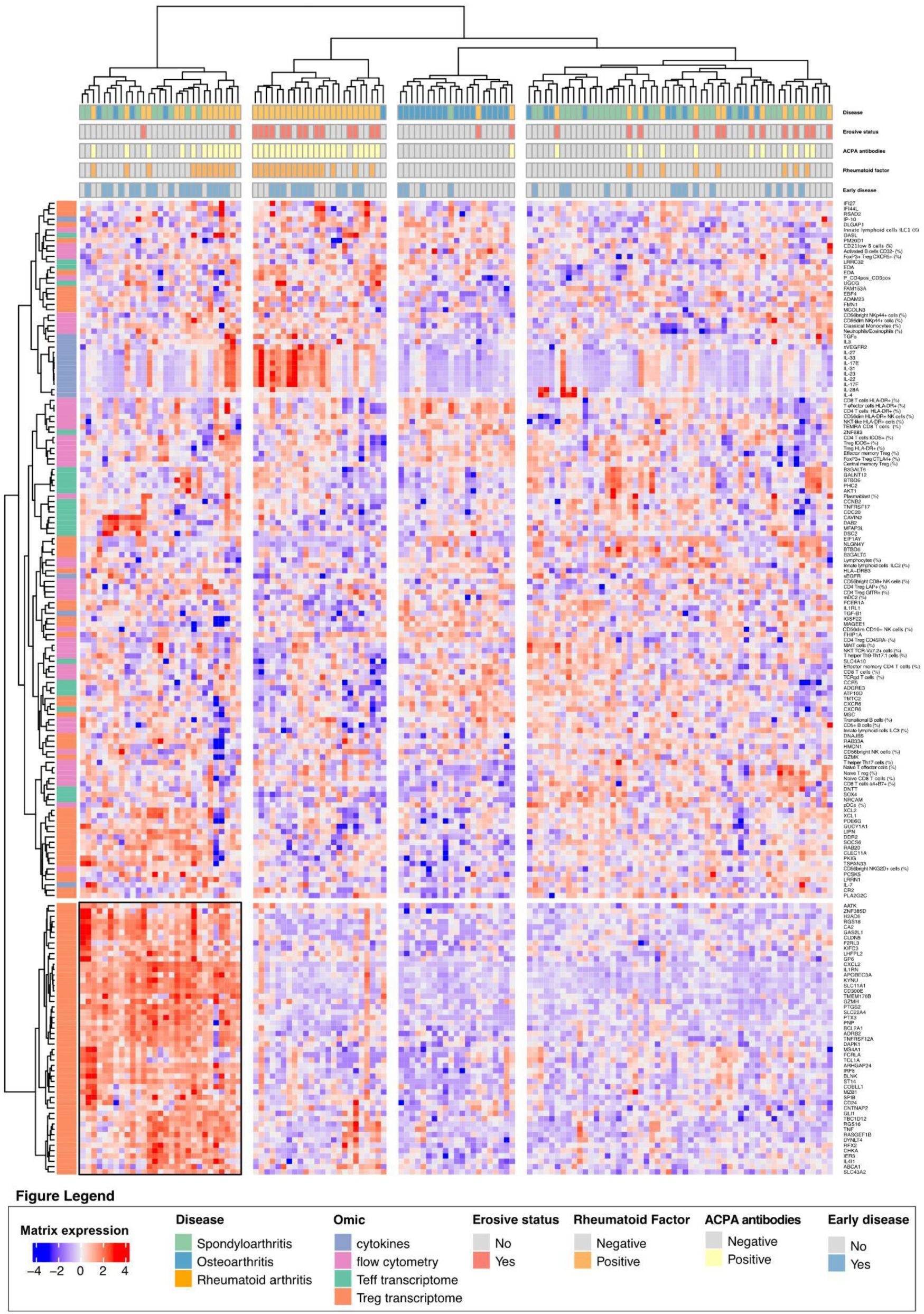
Clustered Image Map (CIM) of the integrative multi-omics signature. CIM heatmap generated with mixOmics from pairwise feature correlations of scaled values. Each row represents an omics feature and each column a patient; rows and columns were ordered by hierarchical clustering. Horizontal side annotations show clinical variables for RA patients, erosive status, rheumatoid factor (RF), anti-citrullinated protein antibodies (ACPA) and early-disease status (defined as disease duration ≤ 2 years), and the vertical side annotation indicates the omics block of origin for each feature. Colour intensity reflects the scaled expression/abundance of each feature.

**Fig. 5.**
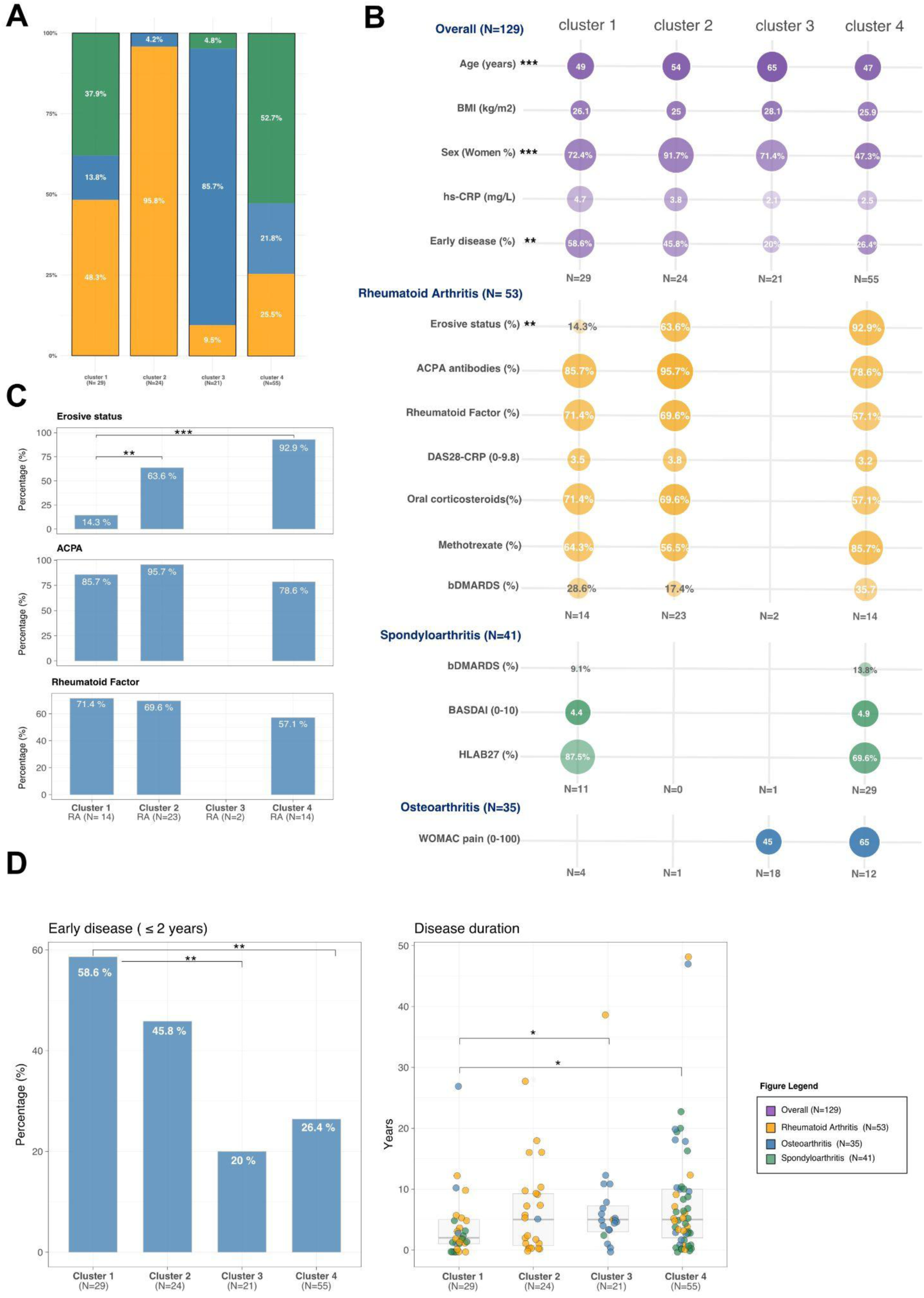
Cluster composition across diseases. **A.** Distribution of clusters within each disease group, expressed as the percentage of patients assigned to each cluster. **B.** Bubble plot of cluster composition within each disease and across the whole cohort (p-values show the Kruskal-Wallis analysis between clusters). **C.** Cluster distribution of erosive status, RF and anti-CCP (ACPA) positivity. **D.** Relationship between erosive status and disease duration across clusters (p-values show the Wilcoxon analysis).

Notably, the composition of cluster 1 was not explained by demographics (age, sex, BMI), treatment (methotrexate, oral corticosteroids or bDMARDs) or disease activity (**Fig 5b-c**). Within RA, patients assigned to cluster 1 showed a markedly lower erosion rate (N=2/14, 14.3%) than those in cluster 2 (N=14/22 63.6%, p=0.0057) or cluster 4 (N=13/14 92.9%, p<0.0001) (Kruskal Wallis p <0.0001), while no significant difference was observed between clusters according to anti-citrullinated protein antibodies (ACPA) and Rheumatoid Factor (RF) status. The principal feature setting cluster 1 apart was early disease (ie ≤ 2 years of disease duration) (N=17/29 58.6%) compared with cluster 2 (N=11/24 45.8%), cluster 3 (N=4/20 20.0% p=0.009) and cluster 4 (N=14/53 26.4% p=0.008) (**Fig. 5c**). When analyzed as a continuous variable, median disease duration was shorter in cluster 1 (median 2.0 years (IQR) (1.0-5.0)) than in cluster 2 (5.0 years (0.8-9.2), ns), cluster 3, (5.0 years (3.0-7.2), p=0.016) and cluster 4, (5.0 years (2.0-10.0), p=0.011) (**Fig. 5d**, **Table S16**). Together, these findings indicate that a multi-omics approach identifies a common, Treg-centered molecular program spanning distinct rheumatic diseases associated with early disease.

### Treg dysfunction as a shared profile in early disease across rheumatic diseases

We next further investigated the identified Treg gene signature comprising 51 genes that were markedly upregulated in cluster 1. Notably, the mean expression of these genes stratified by disease duration revealed a bimodal pattern that distinguished early from late disease (**Fig. 6a**).

**Fig. 6.**
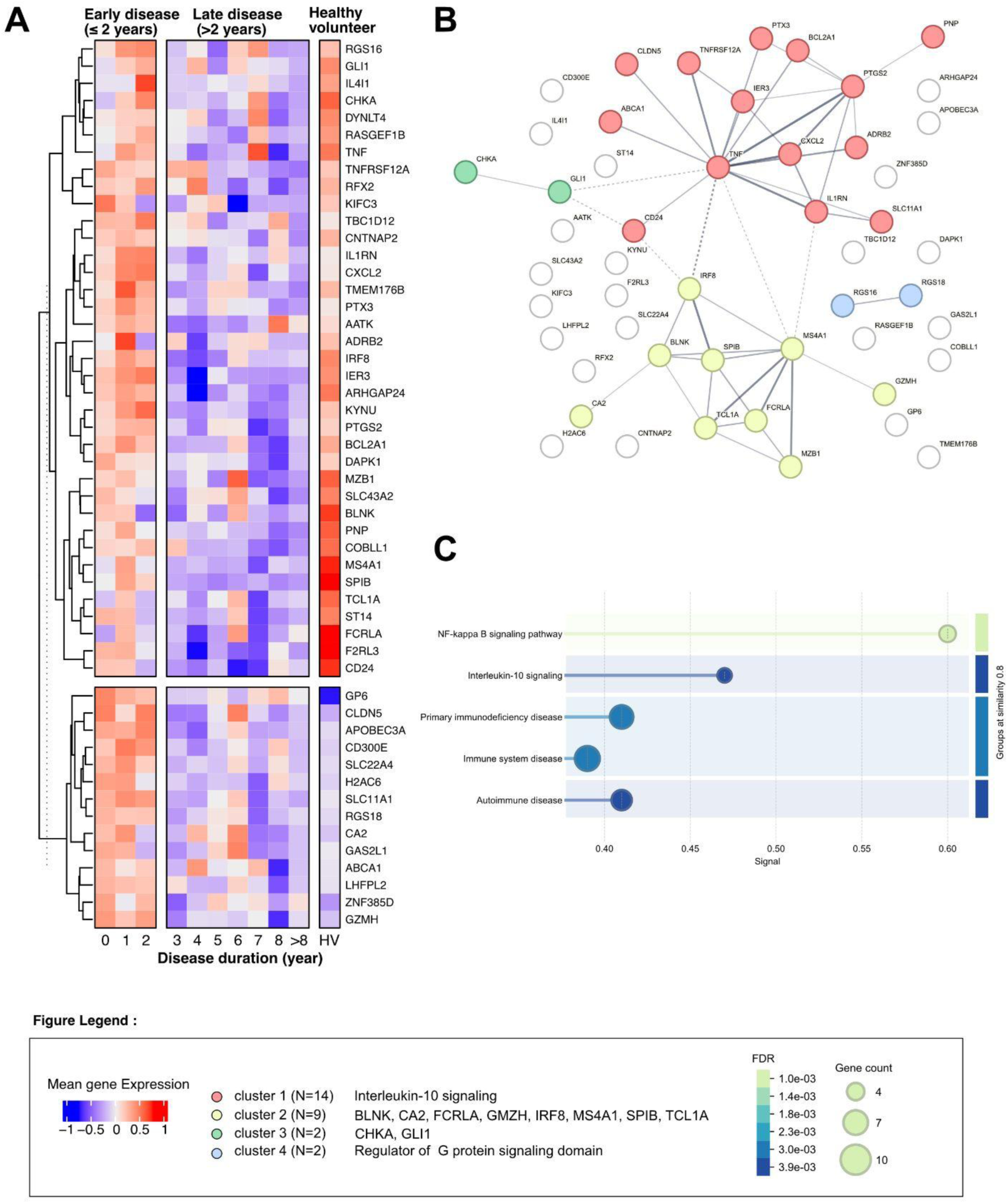
Treg-associated signature with early disease across rheumatic diseases. **A.** Mean expression of the early-disease signature as a function of disease duration, computed on a log₁₀(x + 1)–normalised and scaled expression matrix. **B.** STRING protein–protein interaction network of the signature genes, partitioned into functional modules by the Markov Cluster (MCL) algorithm. **C.** Functional enrichment analysis of the signature (KEGG, DISEASES and Reactome), showing the significantly over-represented pathways.

Mean expression differed significantly between early disease (≤ 2 years) and later stages, defining two opposing profiles. A first set of 37 genes was upregulated in early disease and in HV but downregulated in late disease. Several genes encode proinflammatory mediators or regulators of T cell function, including *TNF*, *TNFRSF12A*, *CXCL2* and *IER3*, alongside genes with immunoregulatory roles such as *IL1RN* (IL-1 receptor antagonist), *IL4I1*, *KYNU* and *IRF8*. A second set of 14 genes was upregulated in early disease yet downregulated in HV, comprising effector and innate-immune genes including *GZMH*, *APOBEC3A*, *CD300E* and *SLC11A1*, together with *SLC22A4*, *RGS18* and the platelet collagen receptor *GP6* and *CA2*. Functional analysis using STRING revealed significant enrichment for genes involved in IL-10 signaling (**Fig. 6b**), together with over-representation of pathways related to NF-κB signaling, primary immunodeficiency, immune system and autoimmune disease (KEGG, Reactome; **Fig. 6c** and **Table S17**). Taken together, these findings indicate that, whereas the multi-omics analysis delineated disease-specific molecular pathways, early disease is characterised by a common Treg-associated dysfunction, marked by an initially intact but subsequently eroded regulatory programme, that may represent a shared pathogenic mechanism across rheumatic conditions evolving over time.

## DISCUSSION

We generated a comprehensive Treg/Teff reference atlas of rheumatic diseases benchmarked against healthy volunteers by combining deep clinical and immune phenotyping across more than 200 well-characterized individuals. The resulting resource integrates over 750 standardized and expert-curated clinical variables with harmonized whole-blood immunophenotyping, multiplex serum cytokine profiling and transcriptomic analysis of purified Treg and Teff populations from matched samples. Each layer was selected to capture complementary aspects of the same immunological process, allowing coordinated cellular, molecular and systemic programmes to emerge. By combining cross-omics integration with sparse feature selection, the analysis identified interpretable molecular signatures that would likely remain obscured if each modality were analyzed independently.

By profiling Tregs and Teffs as distinct, sorted compartments rather than pooled CD4⁺ populations, our data show that the most informative signals -particularly the pronounced, shared transcriptional downregulation within Tregs-would likely be diluted or lost in bulk CD4⁺ analyses. This argues against pooling T-cell subsets with opposing functions and may explain why comparable multi-omics signatures have not previously been reported: to our knowledge, few existing datasets separately have profiled sorted Treg and Teff transcriptomes at this scale, which also limited our ability to pursue external validation.

Integrative multi-omics modelling consistently outperformed individual molecular layers in resolving disease-associated patterns and revealed coordinated molecular programmes that remained undetected through separate analyses. Importantly, this added value arose not simply from combining multiple datasets but from integrating complementary information through a sparse feature-selection framework that identified the most informative variables across molecular layers. Beyond improving model robustness and interpretability, this approach highlights biologically coherent programmes rather than isolated biomarkers, facilitating mechanistic interpretation and downstream validation. These results underscore the added value of integrative modelling, not merely as a technical refinement, but as a means of generating hypotheses that single-omics data cannot support. Unsupervised clustering identified one patient cluster (Cluster 1) that was not driven by disease activity, treatment status or age. Instead, this cluster was characterized by significantly shorter disease duration, being enriched in patients with early disease (≤2 years). Consistent with this, RA patients within the cluster displayed a lower frequency of erosive disease despite comparable antibody positivity, a pattern that independently supports shorter disease duration, given that structural erosions typically require several months to become radiographically apparent^17^. Together, these findings indicate that this cluster captures a shared molecular state associated with early disease rather than disease-specific characteristics. Notably, the Cluster 1 molecular profile was dominated by an altered Treg transcriptomic programme, suggesting that Treg dysfunction may constitute a shared, disease-agnostic feature of the early phase across rheumatic conditions rather than a mechanism confined to any single diagnosis. Hereafter, we use the term “Treg dysfunction” as an operational descriptor of the coordinated cellular and molecular alterations identified in Tregs across rheumatic diseases, without implying experimentally demonstrated impairment of suppressive function.

Within this early-disease signature we identified a dual expression pattern. A first set of genes, upregulated in early disease and in healthy volunteers but lost in later disease, including proinflammatory and T cell–associated genes (*TNF, TNFRSF12A, CXCL2, IER*3) alongside immunoregulatory genes (*IL1RN, IL4I1, KYNU, IRF8*), is compatible with an initially balanced regulatory response that erodes with chronicity. In RA patients, IL-1 receptor antagonists have been described to lower Th17 and Th1 percentages and raise the percentage of Treg cells, with these regulatory changes correlating with clinical improvement^18–20^. A second set, upregulated in early disease but absent in HV (*GZMH, CD300E, APOBEC3A, SLC11A1, SLC22A4, RGS18, GP6, CA2*), included effector, innate-immune and platelet/megakaryocyte-associated genes. The platelet component is of particular interest given the growing recognition of platelets as active participants in immune-mediated inflammatory diseases^21^, and aligns with evidence that platelet-derived mediators can directly impair Treg function^22^. Alongside this, the metabolic (CA2, ABCA1) and chromatin/transcriptional (H2AC6, ZNF385D) genes suggest concurrent metabolic and transcriptional remodelling. Together, this signature is consistent with Tregs responding to inflammatory stress and to the influx of platelet-derived, and other cell-derived signals in early disease.

Functional enrichment centred on IL-10 and NF-κB signalling, immunodeficiency and autoimmune pathways further supports a coherent immunoregulatory axis operating at disease onset. Together these findings argue that Treg dysfunction is not static but dynamic across the disease course, being most pronounced, and potentially most tractable, early on.

These observations carry translational implications. If Treg dysfunction is a shared and time-dependent driver of early rheumatic disease, it defines both a candidate therapeutic target and a specific window in which Treg-directed strategies might be most effective. This aligns with the long-standing concept of a therapeutic “window of opportunity” in RA, typically framed within the first two years of disease, and often the first months, during which immunomodulatory intervention yields disproportionately better structural and clinical outcomes^7,23,24^.Our finding that Treg dysfunction is most pronounced in this same early phase provides a mechanistic rationale for why this window may exist and why it matters. This has direct relevance for the design of future clinical trials, which should account for disease duration and early disease status when stratifying patients and selecting endpoints for Treg-modulating interventions.

Our study has several limitations that also define important directions for future research. While the identified Treg/Teff signatures were derived from a large, deeply phenotyped cohort, validation in independent populations will be important to establish their generalizability. The present cross-sectional design enables the identification of molecular programs associated with different stages of disease but does not directly capture their temporal evolution. Longitudinal studies will therefore be valuable to determine how these programs evolve within individual patients and whether they predict disease progression or therapeutic response. A related consideration is that factors such as age, sex and BMI are known to influence immune cell composition and cytokine production, and may therefore contribute to some of the disease-associated differences observed at the single-omic level, disentangling genuine disease effects from these composite demographic influences will require larger cohorts explicitly powered for stratified or covariate-adjusted analyses. A further consideration relates to the choice of peripheral blood: focusing on this readily accessible, minimally invasive compartment, rather than target tissue such as synovium, allowed us to build a well-powered, multi-layered atlas across three distinct diseases at a scale not achievable with tissue sampling. However, it does not capture site-specific immunopathology, and mechanistic interpretation of the Treg programs identified here should be extended to tissue-level studies. Functional validation of Treg suppressive activity was not performed here; nevertheless, no single assay currently captures the full breadth of Treg suppressive mechanisms, as the classical dose-dependent Teff-proliferation suppression assay principally reflects IL-2 consumption, one of several mechanisms described above. The multi-layered phenotypic and transcriptional signatures reported here may therefore capture a broader picture of Treg dysfunction than any single functional readout, though direct functional confirmation remains an important next step. Finally, extending this integrative framework with functional and single-cell approaches will help resolve the cellular mechanisms underlying the coordinated molecular signatures identified here.

Altogether, this work demonstrates the power of integrative multi-omics analysis to generate hypotheses inaccessible to single-layer approaches, and identifies Treg dysfunction as a candidate shared driver across rheumatic diseases that is dynamic over time. By combining cross-omics integration with interpretable feature selection, our framework resolves coordinated molecular programs that may facilitate biological understanding as well as biomarker discovery. Validating this mechanism prospectively, and testing whether early Treg-directed intervention can alter disease trajectory, represent the logical next steps.

## METHODS

### Patients and Clinical Assessment

The Trans*i*mmunom study (NCT02466217) is a multi-center observational clinical trial designed to cross-phenotype patients with autoimmune and/or autoinflammatory diseases using a multi-omics approach. The comprehensive methodology and procedures have been previously described^25^. Briefly, standardised data collection was ensured through electronic case-report forms (E-CRF) capturing clinical, biological, and radiological data for each participant and 135 mL of peripheral blood were collected from each individual. Enrolment took place from October 2017 to June 2019. The present study focused on three disease cohorts: Rheumatoid Arthritis (RA, n=91), axial and peripheral Spondyloarthritis (SpA, n=58), and knee Osteoarthritis (OA, n=44), alongside healthy volunteers (HV, n=47) classified according to the ACR/EULAR 2010 criteria, the ASAS 2009 criteria, and the ACR criteria for knee OA, respectively^26–28^. For all groups, demographic and clinical data were collected at the time of sampling, including age, sex, body mass index (BMI), and disease duration. Inflammatory status was assessed by high-sensitivity C-reactive protein (hs-CRP) and lymphocyte count. Disease activity was evaluated using validated, disease-specific instruments: the Disease Activity Score 28 based on CRP (DAS28-CRP) for RA, the Bath Ankylosing Spondylitis Disease Activity Index (BASDAI) for SA, and the Western Ontario and McMaster Universities Osteoarthritis Index (WOMAC) pain subscale for OA^29–31^. Radiographic assessment was performed according to standard protocols for each disease. Immunological status was recorded for RA patients, including rheumatoid factor (RF) and anti-citrullinated protein antibody (ACPA) positivity, as well as erosive disease status (in RA patients). HLA-B27 typing was performed where applicable in SpA patients. Treatment data were collected at inclusion, encompassing conventional synthetic disease-modifying antirheumatic drugs (csDMARDs), biological DMARDs (bDMARDs), and oral corticosteroids.

### Flow Cytometry and Deep Immunophenotyping

Blood immunophenotyping was performed using nine flow cytometry panels developed for the Trans*i*mmunom study covering 137 cell populations (**Table S4**)^32,33^. As previously described, Duraclone dry antibody technology was applied to whole blood, which was preferred over peripheral blood mononuclear cells (PBMCs) to minimise technical manipulation and preserve cell phenotype integrity. Panels were designed to characterise B cells, natural killer (NK) cells, monocytes, dendritic cells, mucosal-associated invariant T (MAIT) cells, myeloid-derived suppressor cells, and T cell subsets including activation, migration, memory, polarisation, and regulatory phenotypes. An additional panel incorporating numeration beads enabled determination of absolute cell counts across all populations. All acquisitions were performed on a Gallios cytometer (Beckman Coulter) and analysed with Kaluza software version 1.3 using a common and validated template. For each population, the percentage of parent population, absolute cell count, and mean fluorescence intensity were recorded. Panel composition, cell populations, and gating strategies are detailed in **Figure S1** and **Table S4**. Pairwise comparisons of cell population frequencies between HV and each disease group (RA, SpA, and OA independently) were performed using the two-sided Mann-Whitney U test. P-values were corrected for multiple comparisons across cell populations using the Benjamini-Hochberg procedure, with a FDR-adjusted p-value threshold of 0.05 considered statistically significant.

### Serum Cytokine Quantification

Serum cytokines, chemokines, and soluble receptors were quantified by Luminex® multiplex technology using five MILLIPLEX® kits: Human Cytokine/Chemokine, Human High Sensitivity T Cell, Human Th17, Human TGFβ1, and Human Soluble Cytokine Receptor Magnetic Bead Panel (Merck). Data acquisition was performed on a Luminex® MAGPIX instrument and analysed using xPONENT® software. Instrument calibration and verification were performed one week prior to each experiment. For each kit and batch, two technical controls with expected concentration ranges were duplicated as quality controls. Standards with percentage recovery below 80% or above 120% were excluded from analysis. Analyte concentrations were calculated from mean fluorescence intensity data using a 5-parameter logistic or spline curve-fitting method. All quantified cytokines, chemokines, and soluble receptors are summarized in **Table S5**. Pairwise comparisons between HV and each disease group (RA, SpA, and OA independently) were performed using the two-sided Mann-Whitney U test. P-values were corrected for multiple comparisons across analytes using the Benjamini-Hochberg procedure, with a FDR-adjusted p-value threshold of 0.05 considered statistically significant.

### Regulatory and effector T cell sorting

Peripheral blood (30 mL) was collected for T cell sorting. PBMCs were isolated by Ficoll-Paque density gradient centrifugation. CD4+ T cells were enriched using EasySep™ magnetic beads and subsequently sorted into regulatory T cells (Treg: CD25+CD127loCD4+) and effector T cells (Teff: CD25-CD127hiCD4+) according to the manufacturer’s recommendations. Sort purity was confirmed by flow cytometry using anti-CD4 and anti-FoxP3 antibodies; only samples with a minimum purity threshold of 80% were retained for downstream experiments. Sorting purity ranged from 92% to 98% for Teff cells and from 85% to 98% for Treg cells.

### mRNA Transcriptomic profiling and preprocessing

Total RNA was extracted from sorted Treg and Teff cells using the RNAqueous™-Micro Total RNA Isolation Kit (Invitrogen, Waltham, MA) following the manufacturer’s instructions. RNA quality was assessed using either the Bioanalyzer 2100 or TapeStation 4200 (Agilent), with a RNA Integrity Number (RIN) threshold of ≥7 applied as an inclusion criterion (input: 10 ng for Tregs, 100 ng for Teff). Isolated RNA was reverse-transcribed using an Oligo(dT) primer, and cDNA was amplified, fragmented, and tagged using the SMARTer® Ultra® Low Input RNA Kit for Sequencing v4 (Takara Bio USA, Mountain View, CA) by the iGenSeq core facility (Paris Brain Institute). RNA sequencing was performed on an Illumina NextSeq 500 platform (75 bp, paired-end) targeting an average depth of 15 million reads per sample by the iGenSeq core facility (Paris Brain Institute). Raw reads were trimmed using Trim Galore and pseudo-aligned to the GRCh38.p13 Ensembl human transcriptome reference using Salmon 2.0, with transcript-level abundances imported into R using the tximport package (v1.18.0)^34,35^.

### Differential gene expression analysis and functional enrichment analysis

Differential gene expression between each disease group (RA, SpA, OA) and HV was performed separately for Treg and Teff populations using DESeq2 (v1.38.3)^36^. Quality control was performed at both sample and gene levels. Genes with counts below 20 in fewer than 20 patients were filtered out from the normalised count matrix, and genes with a base mean below 8 were additionally excluded prior to testing. Quality control metrics including variance stabilisation transformation, Cook’s distance, and dispersion estimates were applied. Potential confounding variables including sex, BMI, age, sequencing lane, and batch were not included as model covariates but were visually assessed through principal component analysis and sample-level quality control plots to evaluate their contribution to transcriptomic variance, as summarised in **Fig S3-S5**. Multiple testing correction was applied using the Benjamini-Hochberg procedure; genes with an FDR-adjusted p-value below 0.05 and an absolute log2 fold change exceeding log2(2) (i.e. >2-fold change) were considered differentially expressed.

Over-representation analysis (ORA) was performed using the clusterProfiler R package^37^ against Gene Ontology Biological Process (GO-BP) terms. For each disease group and cell type, nominally significant genes (raw p-value ≤ 0.05) were stratified by direction of effect (upregulated (log2FC > 0) and downregulated (log2FC < 0)), and tested independently to preserve biological directionality. Background universe comprised all genes detected and retained after low-count filtering in the respective DESeq2 model, converted to Entrez IDs via the org.Hs.eg.db annotation package. Enrichment testing was performed with a nominal p-value threshold of 0.05, Benjamini-Hochberg multiple testing correction, and a q-value threshold of 0.2. Redundant GO terms were subsequently removed using semantic similarity-based simplification with a similarity cutoff of 0.7, retaining the term with the minimum adjusted p-value within each redundant cluster, as implemented in the simplify function of clusterProfiler. Enriched terms with an adjusted p-value below 0.05, a gene ratio exceeding 0.03, and a minimum of 20 annotated genes were retained for visualization. All analyses were conducted in R version 4.2.2 (2022-10-31).

### Multi-omics integration by multi-block sPLS-DA

Multi-omics integration was performed with multi-block sparse partial least-squares discriminant analysis (sPLS-DA) in the mixOmics R package (v6.23.4)^38–40^, restricted to the 129 patients with complete data across all four blocks (flow cytometry, cytokines, Treg transcriptome, Teff transcriptome). Multi-block sPLS-DA is a supervised extension of sparse generalized canonical correlation analysis (sGCCA) that generalizes PLS and PLS-DA to multiple matched datasets^41,42^. It seeks latent components (linear combinations of the variables within each block) such that the sum of pairwise covariances between blocks and the outcome is maximized, with each block pair weighted by a user-specified design matrix. Sparsity is imposed by lasso (L1) penalization, retaining a defined number of variables per block on each component to yield an interpretable discriminant signature.

Each block was log₁₀(x + 1)–transformed and scaled prior to integration. Blocks were combined with the centroid scheme (SVD initialization), and models were fitted to discriminate the three disease groups (OA, SpA, RA). We used a null design matrix (off-diagonal weight = 0), prioritizing class discrimination over the recovery of cross-block correlation. Two components were retained, consistent with the K − 1 rule for three classes and confirmed by the weighted-vote error-rate criterion. The number of variables selected per block and component was tuned by exhaustive search over the full grid of 1,250 combinations using repeated 5-fold cross-validation.

Classification performance was estimated by repeated five-fold cross-validation and reported as the mean and standard deviation of the error rate across repeats. Averaged prediction, weighting all blocks equally, was used for performance evaluation and for benchmarking against single-block models, because equal weighting permits a like-for-like decomposition into per-block error rates. Weighted prediction, weighting each block by the correlation of its component with the outcome so that blocks more strongly associated with disease contributed more to the final call, was used for hyperparameter tuning, with the balanced error rate (BER) as the optimization measure to limit class-imbalance bias; because this weighting is defined only once the blocks are combined, the weighted error rate exists only at the consensus level and cannot be decomposed by block. The weighted consensus error rate on the final two-component model is reported as a sensitivity analysis. Discrimination of each disease from the others was additionally summarized by one-versus-rest area under the ROC curve (AUC) on the second component. Component loadings were inspected to identify the variables driving discrimination across diseases.

Multi-omics variables were integrated and explored through a correlation-based clustered image map (CIM). Pairwise associations between variables were quantified using Pearson correlation coefficients. The resulting matrix was converted into a distance matrix defined as 1 − r, and unsupervised hierarchical clustering was performed on these distances using Ward’s minimum-variance method (Ward.D2 agglomeration). The clustered correlation structure was rendered as a heatmap using the ComplexHeatmap R package (2.14.0), with dendrograms ordering both rows and columns according to the hierarchical clustering solution.

### Cluster characterization, early disease and functional enrichment analysis

Cluster composition was subsequently characterized with respect to demographic and clinical variables. Variables assessed across all patients included sex, age, body mass index (BMI), early disease status (defined as disease duration ≤ 2 years) and high-sensitivity C-reactive protein (hs-CRP). Disease-specific variables were additionally evaluated. For RA, these comprised immunological status (anti-citrullinated protein antibodies [ACPA] and rheumatoid factor [RF]), disease activity (DAS28-CRP), and treatment with methotrexate, biologic disease-modifying antirheumatic drugs (bDMARDs) and oral corticosteroids (defined as oral prednisone intake ≥ 5 mg/day). For SpA, variables comprised HLA-B27 status, disease activity (BASDAI) and bDMARD treatment. For OA, the WOMAC index and associated pain scores were assessed. Between-cluster differences in clinical variables were evaluated using non-parametric methods. Continuous variables were compared with the Mann–Whitney U test (equivalent to the Wilcoxon rank-sum test), applied pairwise between clusters as a post hoc analysis. Categorical variables were compared using the chi-square or Fisher’s exact test, as appropriate (p < 0.05).

Functional analysis of the early-disease signature was performed using STRING DB. Mean expression of Treg genes were computed per patients grouped by year of disease duration as the average scaled expression across signature genes, on a log₁₀(x + 1)–normalised and z-scaled expression matrix and plotted as a function of disease duration (years). Over-representation analysis of the early-disease signature genes was performed using STRING against the DISEASES, KEGG and Reactome annotation databases. For each enriched term, the table reports the category, the term identifier and description, the number of observed versus background genes, the enrichment strength (log₁₀ of observed/expected), the signal score, the false discovery rate (Benjamini–Hochberg corrected), and the matching proteins in the network (Ensembl protein IDs and gene labels). Only terms passing a false discovery rate < 0.05 are shown. Enriched terms included immune system disease, primary immunodeficiency disease and autoimmune disease (DISEASES), NF-κB signalling (KEGG), and interleukin-10 signalling (Reactome). Protein–protein interaction network was partitioned into functional modules using the Markov Cluster (MCL) algorithm with an inflation parameter of 3, and modules were annotated according to their enriched functional terms.

## ACKNOWLEDGMENTS

We thank Dr Karine Louati, Dr Ariane Do, Dr Sandra Desouches, Dr Sabine Trellu and Dr Julien Champey from the Rheumatology Department of Saint Antoine for help on patient sampling as well as Ms Cornelia Degbe and Nathalie Ferry from the Biotherapy Department of Pitié Salpêtrière hospital for sample handling. We thank the members of the URC Lariboisière for their support in the eCRF data record and validation. This work benefited from equipment and services from the iGenSeq core facility at the Paris Brain Institute (ICM, Paris, France) for all the RNA-seq data production.

## COMPETITINGS INTEREST

JS received consulting or conference fees from Grünenthal, Pfizer, Abbvie, Janssen, Novartis, AlfaSigma, Abbvie, Lilly, UCB, Fresenius Kabi, Guerbet, Axomove, IBSA and Nordic Pharma.

## AUTHORS CONTRIBUTION

DK conceptualized the whole study, designed, received funding, and coordinated the Trans*i*mmunom LabEx project as well as the Trans*i*mmunom trial, and contributed to all analyses. MB, JS and EMF designed the multi-omics analytical framework. MB performed the integrative analyses. JS, FB, CR, RL and AC recruited participants and collected clinical data. EMF, MB, MR, MS, AR, FP, AS, PS, HV, VD, NC, LA, PB, VM and NT generated and processed the molecular and immunological datasets. FM and PH contributed mathematical expertise for data analysis. CA coordinated Trans*i*mmunom project management. MB drafted the manuscript, with critical input from all authors.

## DATA AND CODE AVAILABILITY

This manuscript is a preprint and has not yet undergone peer review. The data and code supporting the findings of this study will be made publicly available following peer-reviewed publication. During peer review, all datasets underlying the reported results, including any not yet publicly released, will be made available to editors and reviewers on request.

