## Supplementary material for "A regulatory T cell signature provides a shared molecular basis for the therapeutic window of opportunity in rheumatic disease": Multi-omics TI Supplemental.docx

**Supplementary Figures**

Fig. S1 | Deep immunophenotyping gating strategy adapted from Pitoiset et al.

Fig. S2 | Scatter plot of deep Immune Phenotyping across rheumatologic diseases.

Fig. S3 | Transcriptomic quality control for Teff cells. Quality control metrics for CD4+ effector T cells (Teff).

Fig. S4 | Transcriptomic quality control for Treg cells. Quality control metrics for CD4+ regulatory T cells (Treg).

Fig. S5 | Principal component analysis of Teff and Treg transcriptomes.

Fig. S6 | Volcano plots of differential gene expression in Teff cells.

Fig. S7 | Heatmap of differential expression significance and mean group expression in Teff cells.

Fig. S8 | Volcano plots of differential gene expression in Treg cells.

Fig. S9 | Heatmap of differential expression significance and mean group expression in Treg cells.

Fig. S10 | UpSet plot of shared and unique features across diseases.

Fig. S11 | Cross-validated classification error rate as a function of the number of components and prediction distance.

Fig. S12 | Overall, per-disease and per-block classification error rates.

Fig. S13 | Per-block discrimination of each disease by ROC analysis.

Fig. S14 | Side heatmap of the multi-omics analysis with clinical variables.

Fig S15 | Cluster composition for age and sex

**Supplementary Tables**

Table S1 | Inclusion and Exclusion criteria of the Trans*i*mmunom study

Table S2 | Rheumatoid arthritis clinical characteristic according to disease activity (N=91)

Table S3 | Spondyloarthritis clinical characteristic according to disease activity (N=58)

Table S4 | Deep immunophenotyping cells populations and panels.

Table S5 | Cytokines and Panel measured in the Trans*i*mmunom study.

Table S6 | Cell populations associated with rheumatologic diseases (N = 36).

Table S7 | Cytokines associated with rheumatologic diseases (n=47).

Table S8 | Differentially expressed genes in Teff across disease groups versus HV (n=177)

Table S9 | Differentially expressed genes in Treg across disease groups versus HV (n=262)

Table S10 | Over-representation analysis of Gene Ontology Biological Process terms in Teff and Treg cells across disease groups.

Table S11 | Discriminatory performance (AUC) of immunological measurement modalities for rheumatic diseases.

Table S12 | Multi-omics feature loadings in component 1 discriminating osteoarthritis, spondyloarthritis, and rheumatoid arthritis.

Table S13 | Multi-omics feature loadings in component 2 discriminating osteoarthritis, spondyloarthritis, and rheumatoid arthritis.

Table S14 | Composition of the four clusters according to clinical and biological characteristics, overall and by disease group

Table S15 | Post-hoc pairwise comparisons between clusters for variables showing a significant overall difference.

Table S16 | Disease duration across clusters: additional comparison supporting the early-disease finding.

Table S17 | Markov Cluster (MCL) algorithm partitioning of the early-disease signature protein–protein interaction network.

Table S18 | Functional enrichment analysis of the early-disease signature.

**Fig. S1 | Deep immunophenotyping gating strategy, adapted from Pitoiset et al.** Panels and targeted cell populations are highlighted in color. Cell populations included in the enumeration panel, which allows bead-based counting and the identification and quantification of all populations, are highlighted in blue.

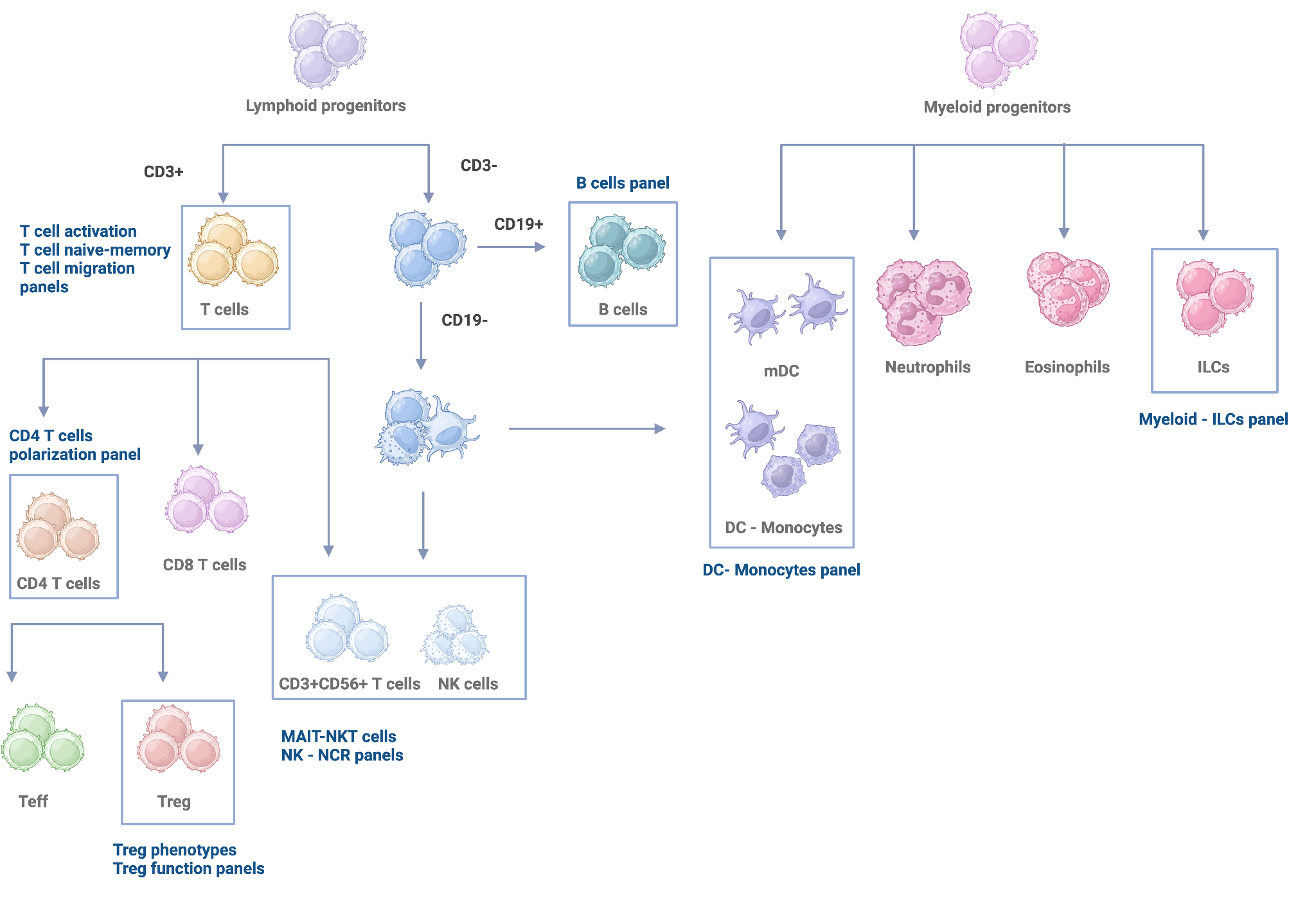

CD, Cluster of differentiation; ILC, Innate lymphoid cells; MAIT, MucoSpAl-associated invariant T cells; NK, Natural Killer, ILC, Innate lymphoid cells; NK, Natural killer; NKT, Natural Killer T cells; Teff T effectory; Treg, T regulatory

**Fig. S2 | Scatter plots of deep immunophenotyping across rheumatologic diseases.** Scatter plots show cell population percentages, as determined by flow cytometry, across the Transimmunom cohort (N = 240): HV (N = 47), OA (N = 44), RA patients (N = 91), and SpA patients (N = 58). Differences between each disease group and the HV reference group were assessed using the non-parametric Wilcoxon rank-sum test (Mann-Whitney U test), with P values adjusted for multiple testing by the Benjamini-Hochberg procedure. Only cell populations showing significant differences (adjusted P ≤ 0.05) are shown.

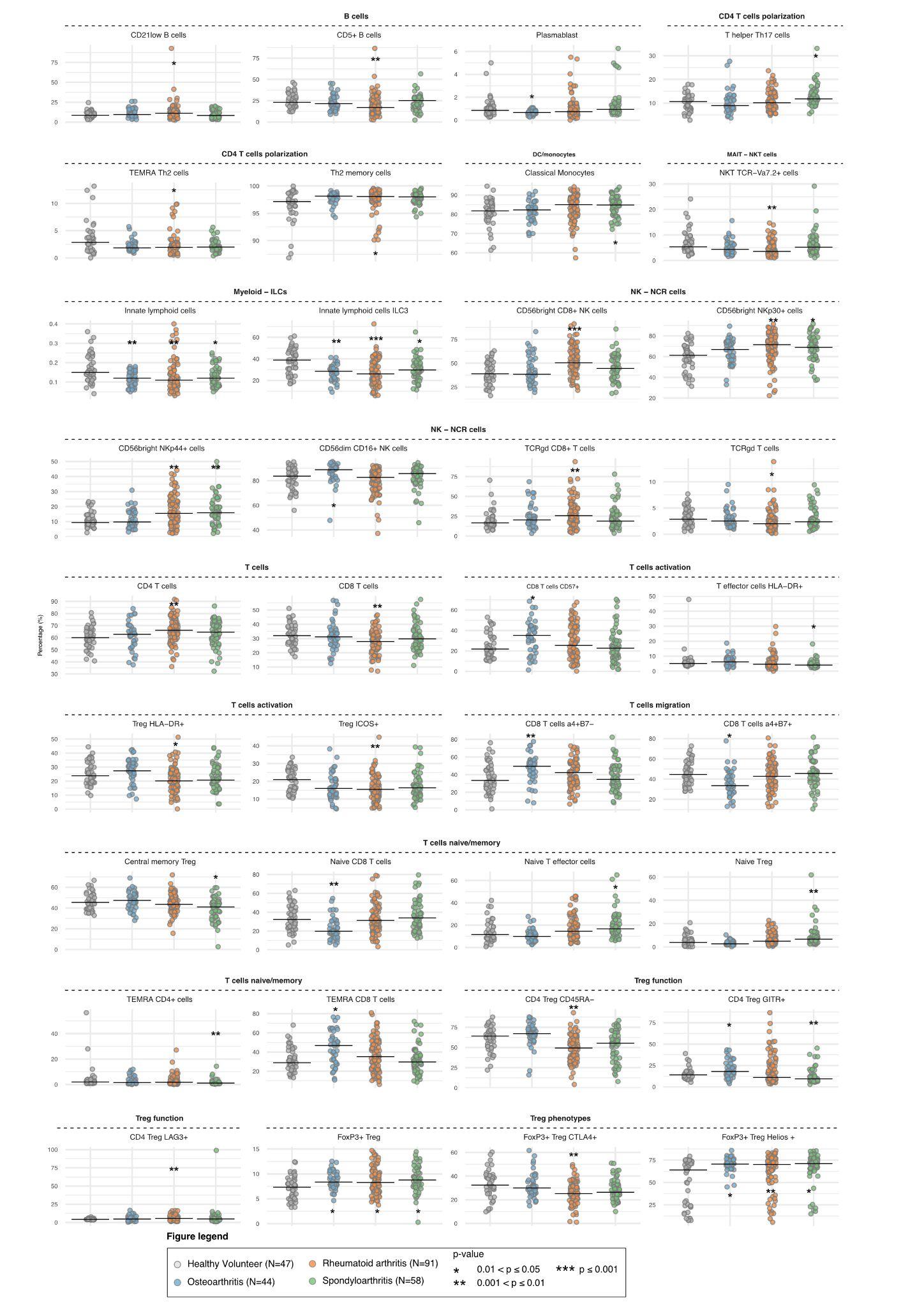

**Fig. S3 | Transcriptomic quality control for Teff cells. Quality control metrics for CD4+ effector T cells (Teff).** A Variance stabilising transformation (VST) of normalised counts across samples. B. Cook's distance per sample, used to identify outliers with disproportionate influence on differential expression estimates. C. Principal component analysis (PCA) of VST-normalised expression profiles, with samples coloured by cell type, sex, batch, disease, BMI age to assess the contribution of technical and biological variables to transcriptomic variance.

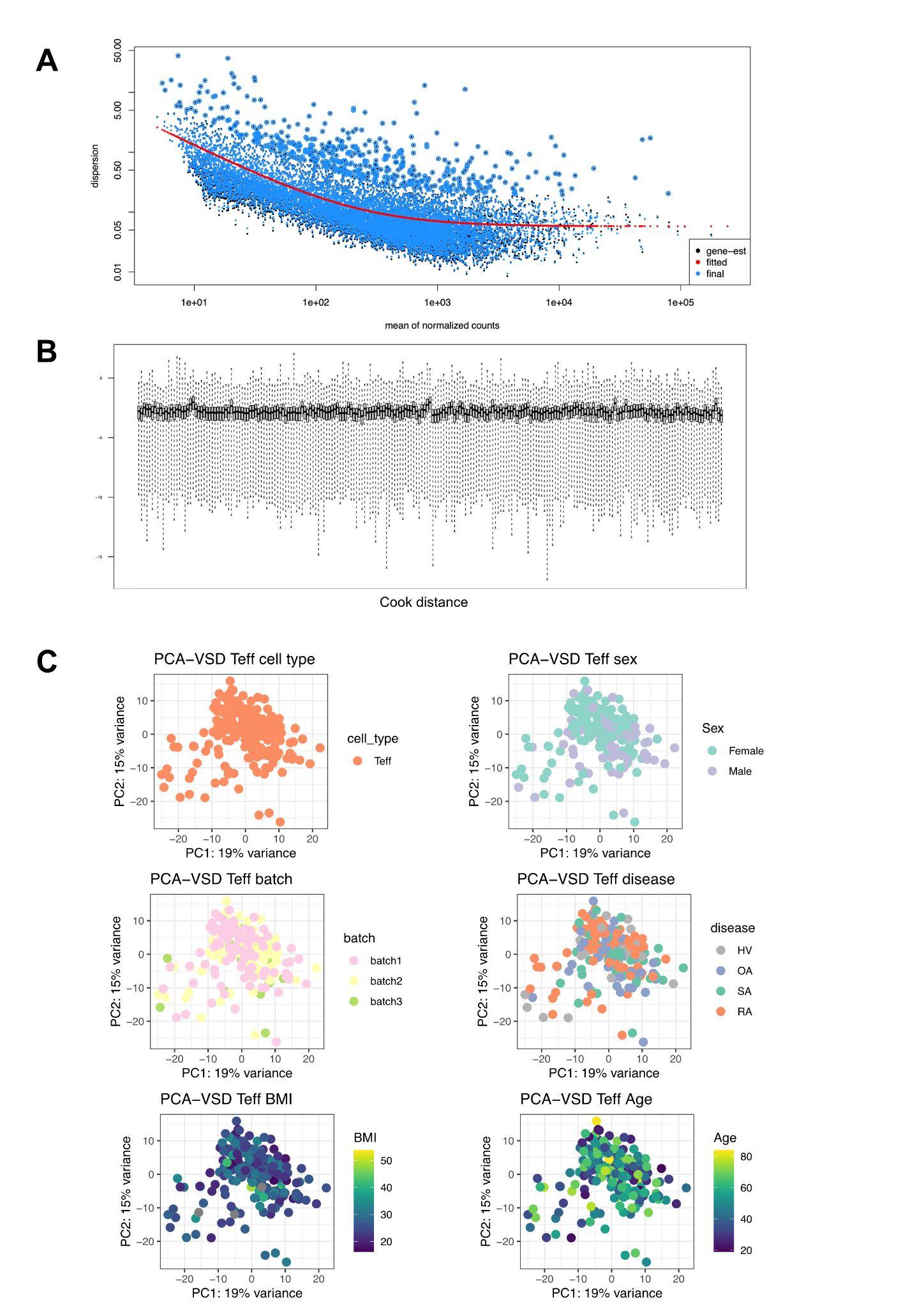

**Fig. S4 | Transcriptomic quality control for Treg cells. Quality control metrics for CD4+ regulatory T cells (Treg).** A Variance stabilising transformation (VST) of normalised counts across samples. B. Cook's distance per sample, used to identify outliers with disproportionate influence on differential expression estimates. C. Principal component analysis (PCA) of VST-normalised expression profiles, with samples coloured by cell type, sex, batch, disease, BMI age to assess the contribution of technical and biological variables to transcriptomic variance.

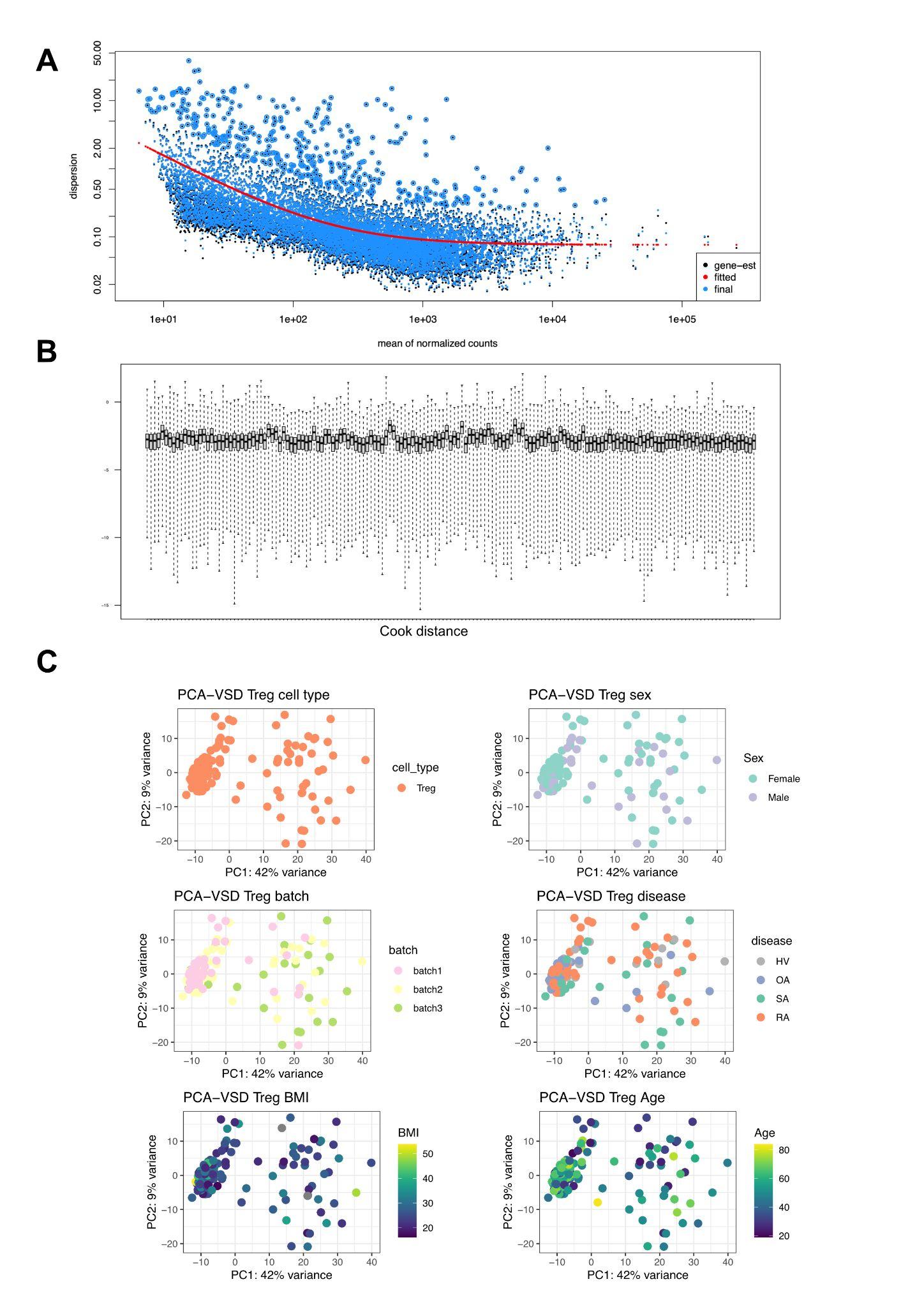

**Fig. S5 | Principal component analysis of Teff and Treg transcriptomes.** Joint PCA projection of VST-normalised gene expression profiles from CD4+ effector T cells (Teff) and CD4+ regulatory T cells (Treg), illustrating the transcriptomic separation between cell types and disease groups. Each point represents one sample, coloured by disease group (RA, SpA, OA, HV) and shaped by cell type.

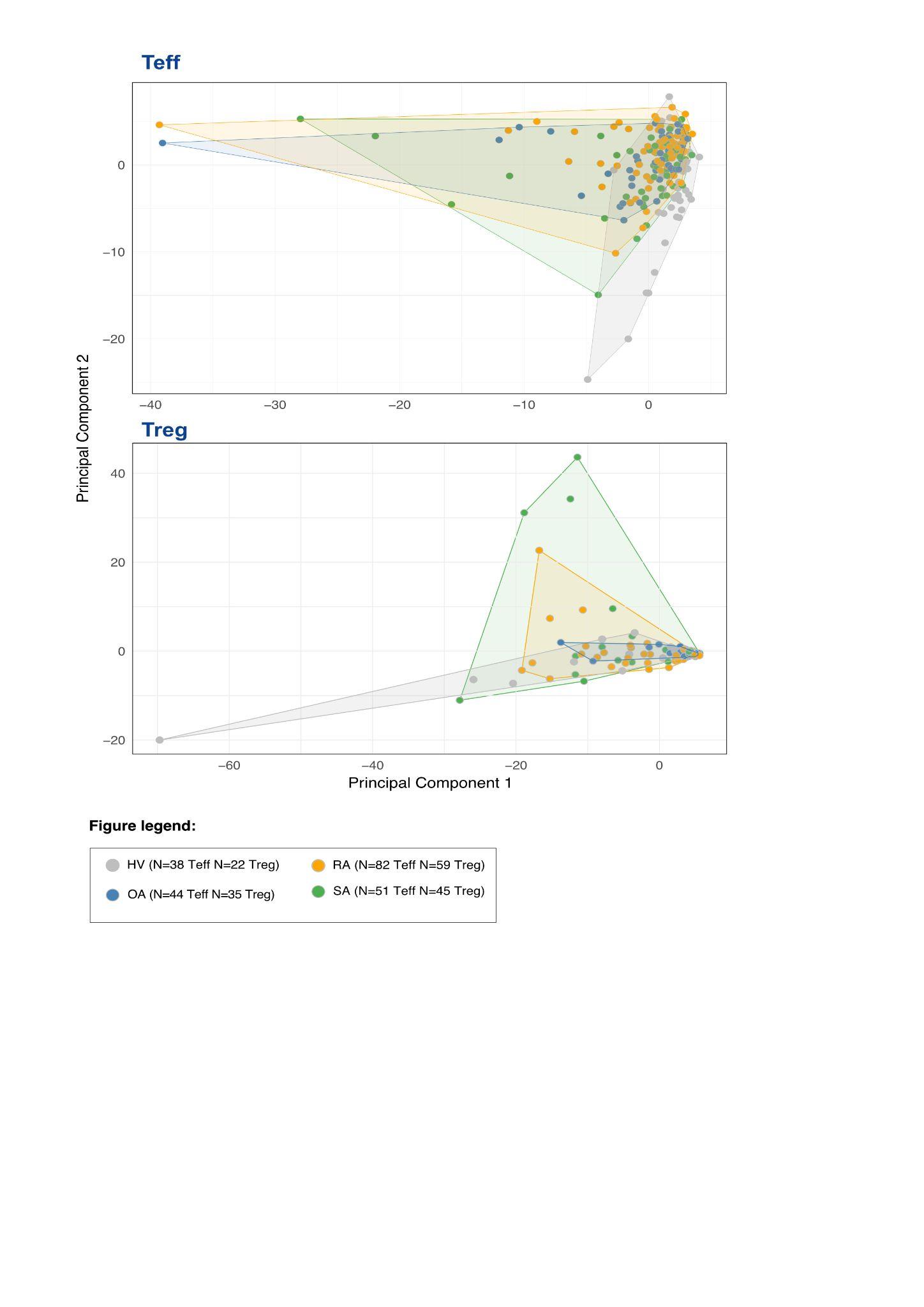

**Fig. S6 | Volcano plots of differential gene expression in Teff cells.** Differential gene expression results for CD4+ effector T cells (Teff) in each disease group versus healthy volunteers (RA vs HV, SpA vs HV, OA vs HV). Each point represents one gene; the x-axis shows log2 fold change and the y-axis shows −log10 Benjamini-Hochberg adjusted p-value. Genes meeting the significance threshold (FDR < 0.05, |log2FC| > log2(2)) are highlighted, with upregulated genes shown in red and downregulated genes in blue. The total number of significant genes per comparison is annotated.

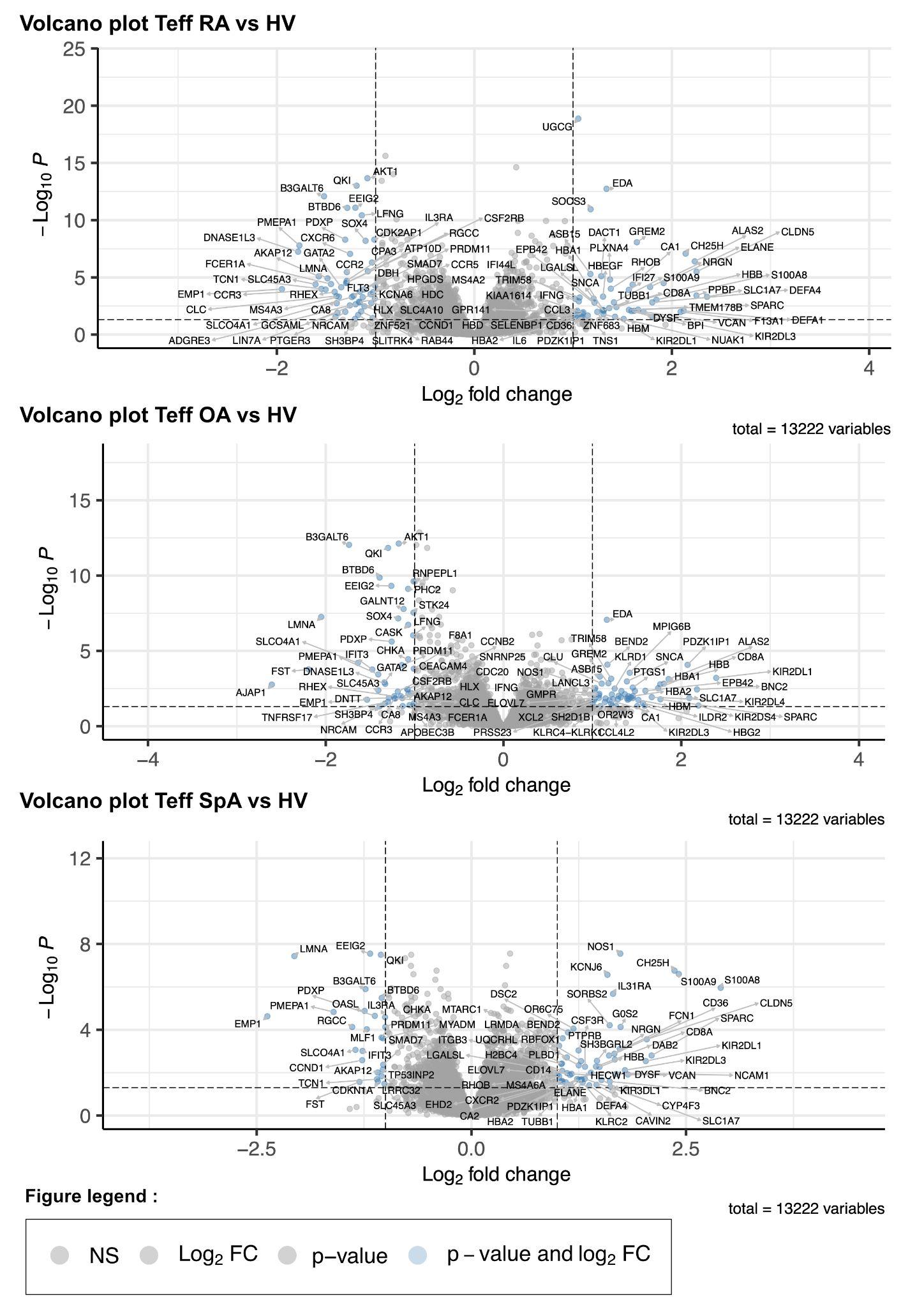

**Fig. S7 | Heatmap of differential expression significance and mean group expression in Teff cells.** Heatmap displaying nominal p-values and mean normalised expression per disease group for differentially expressed genes identified in CD4+ effector T cells (Teff). Rows represent genes and columns represent disease groups (RA, SpA, OA, HV). Colour intensity reflects mean normalised expression; significance of pairwise comparisons versus healthy volunteers is indicated by a dot (FDR ≤ 0.05)

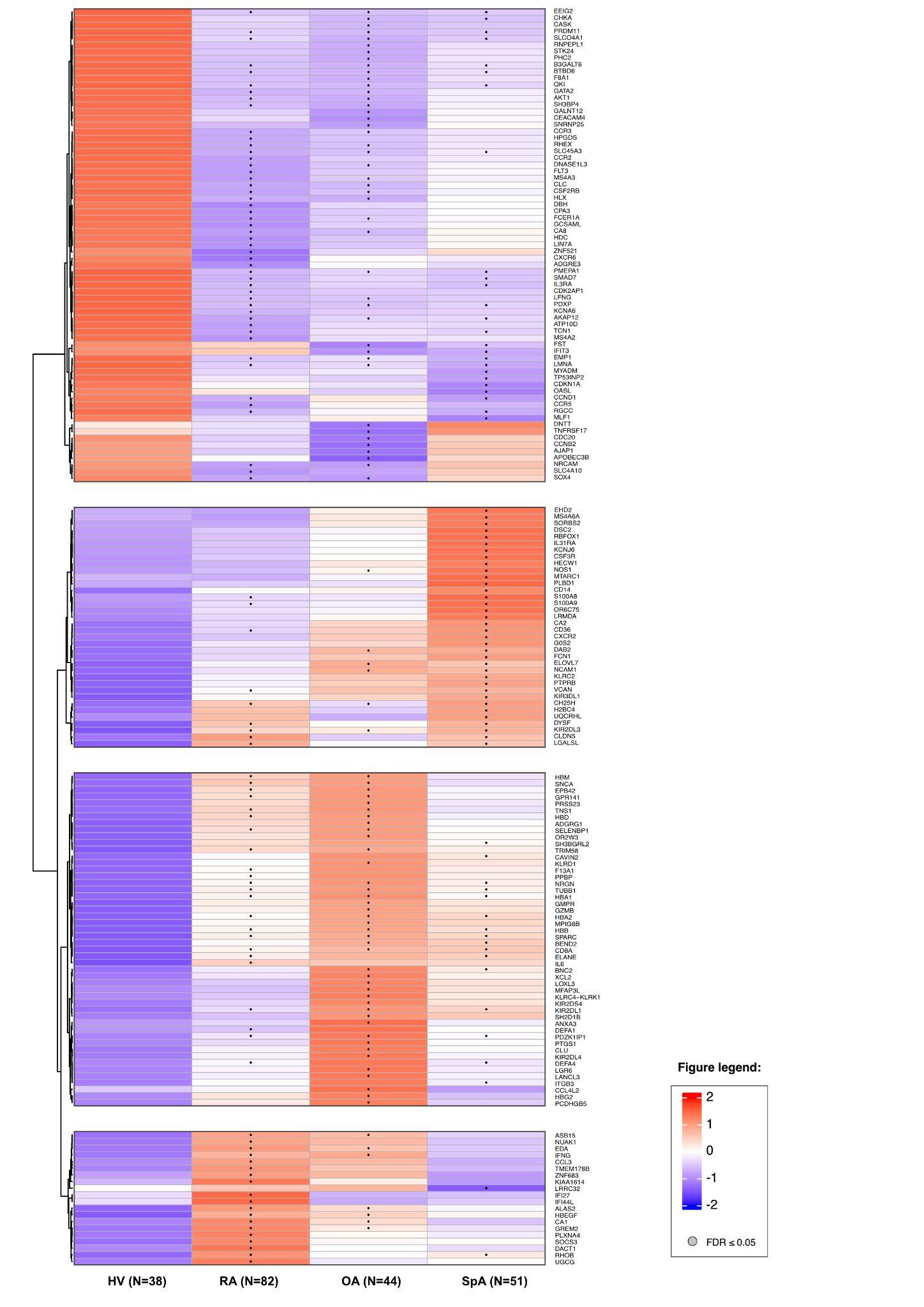

**Fig. S8 | Volcano plots of differential gene expression in Treg cells.** Differential gene expression results for CD4+ regulatory T cells (Treg) in each disease group versus healthy volunteers (RA vs HV, SpA vs HV, OA vs HV). Each point represents one gene; the x-axis shows log2 fold change and the y-axis shows −log10 Benjamini-Hochberg adjusted p-value. Genes meeting the significance threshold (FDR < 0.05, |log2FC| > log2(2)) are highlighted, with upregulated genes shown in red and downregulated genes in blue. The total number of significant genes per comparison is annotated.

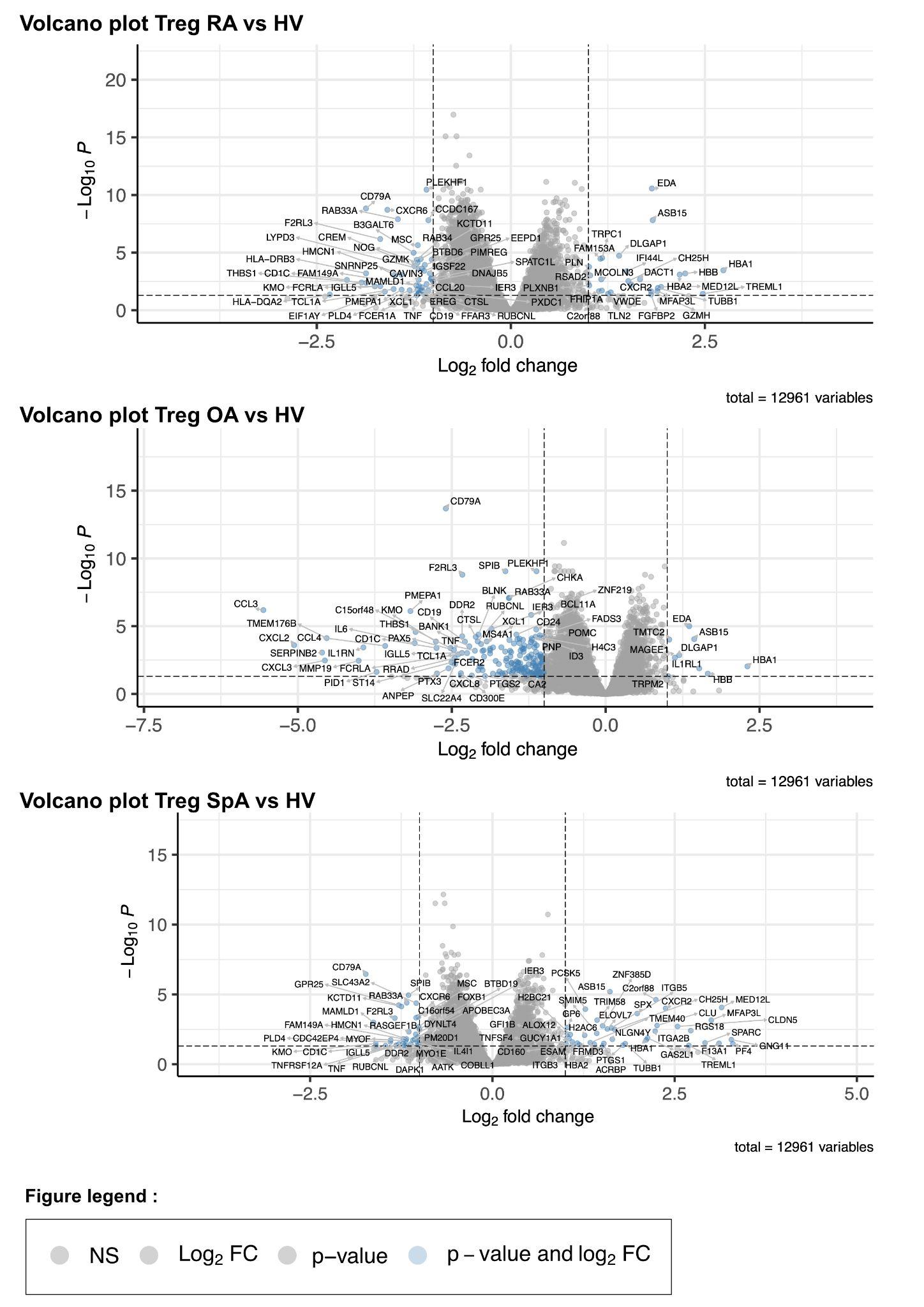

**Fig. S9 | Heatmap of differential expression significance and mean group expression in Treg cells.** Heatmap displaying nominal p-values and mean normalised expression per disease group for differentially expressed genes identified in CD4+ regulatory T cells (Treg). Rows represent genes and columns represent disease groups (RA, SpA, OA, HV). Colour intensity reflects mean normalised expression; significance of pairwise comparisons versus healthy volunteers is indicated by a dot (FDR ≤0.05).
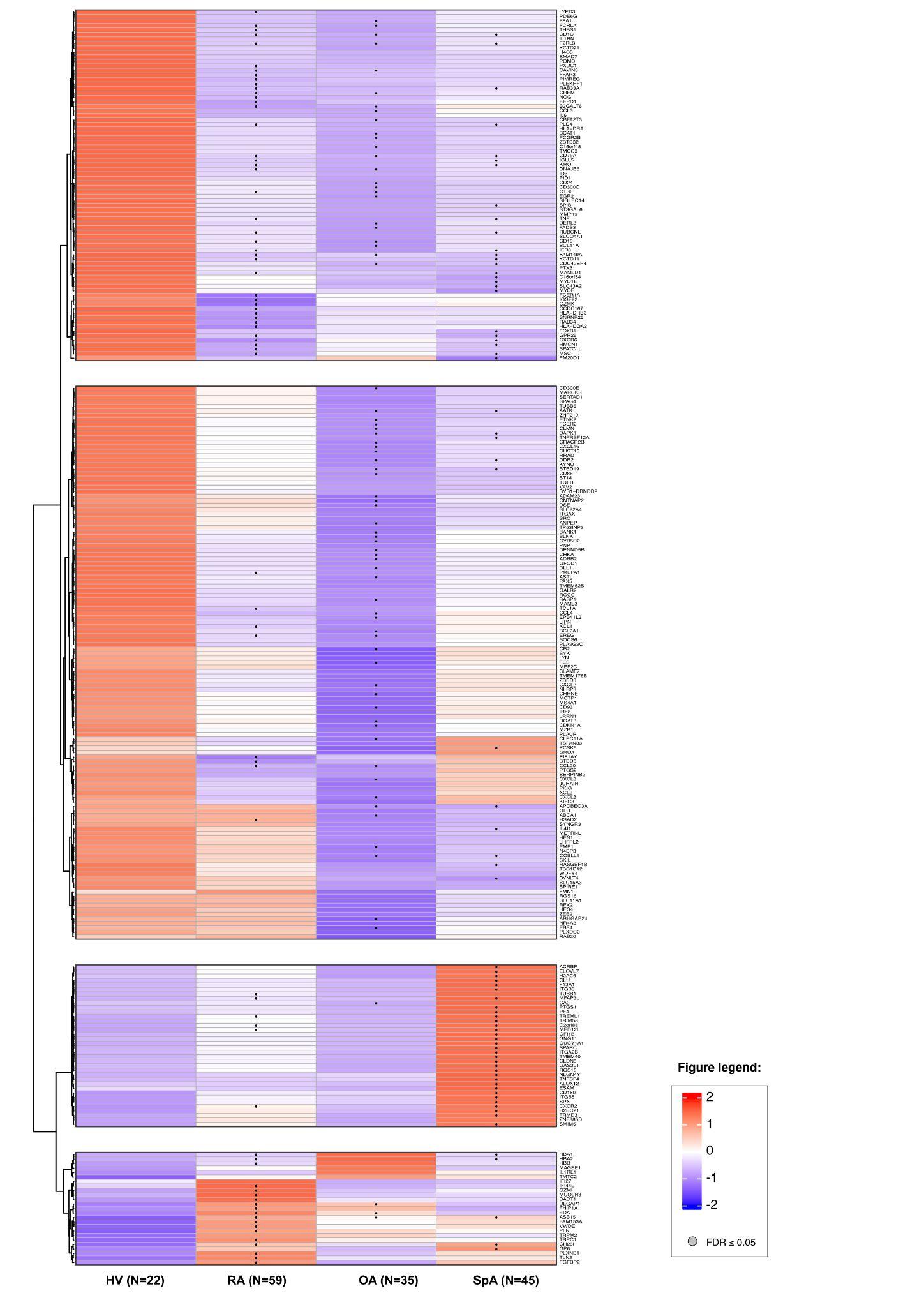

**Fig. S10 | UpSet plot of shared and unique features across diseases.** UpSet plot showing the number of differentially abundant features that are unique to, or shared between, osteoarthritis (OA), spondyloarthritis (SpA) and rheumatoid arthritis (RA), within each omics layer (flow cytometry, cytokines, Treg transcriptome and Teff transcriptome).

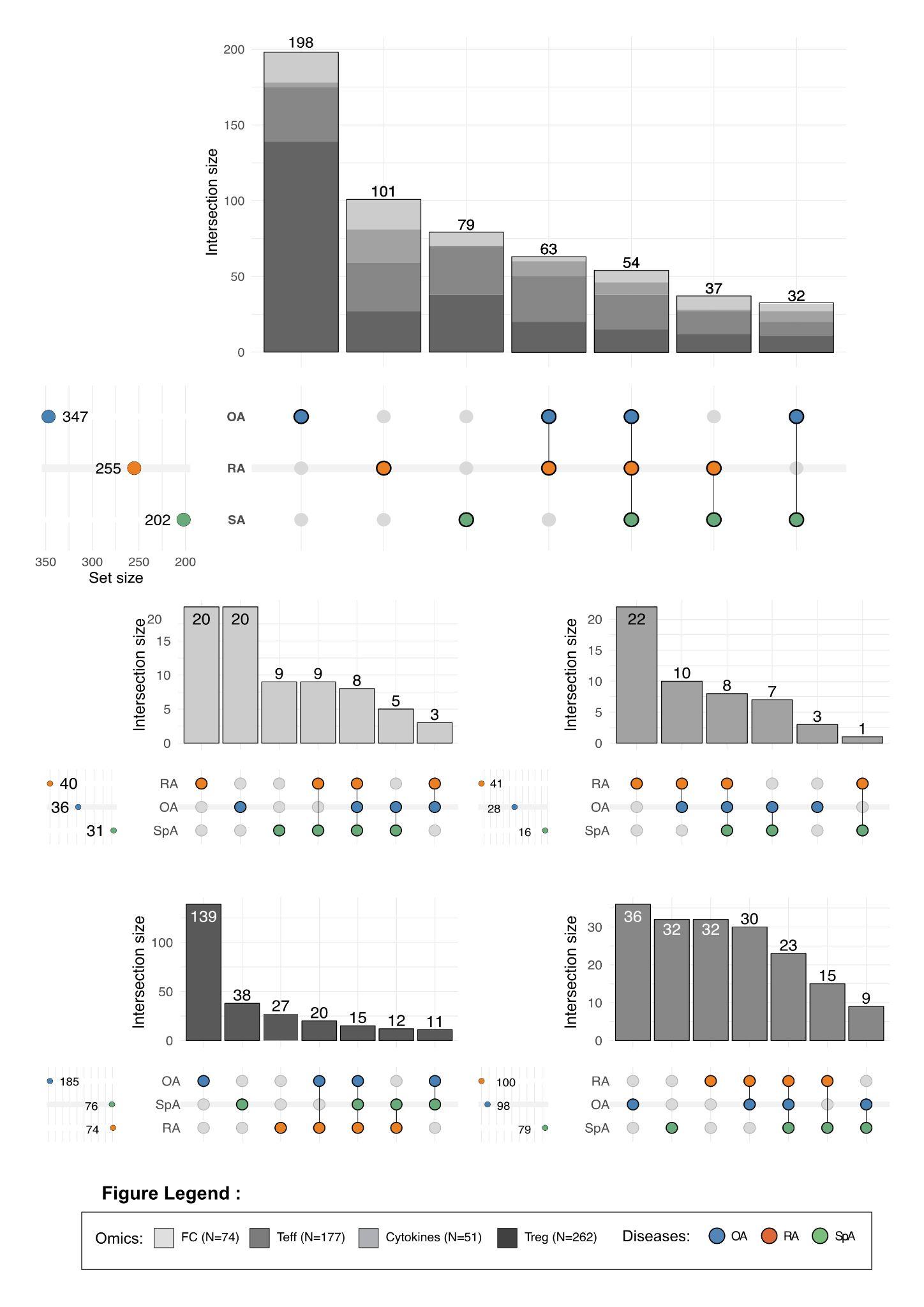

**Fig. S11 | Cross-validated classification error rate as a function of the number of components and prediction distance.** Overall error rate (ER) and balanced error rate (BER) of the integrated multi-block sPLS-DA model estimated by repeated five-fold cross-validation and plotted against components 1 and 2. Error rates are shown separately for the three prediction distances (maximum distance, centroid distance and Mahalanobis distance).

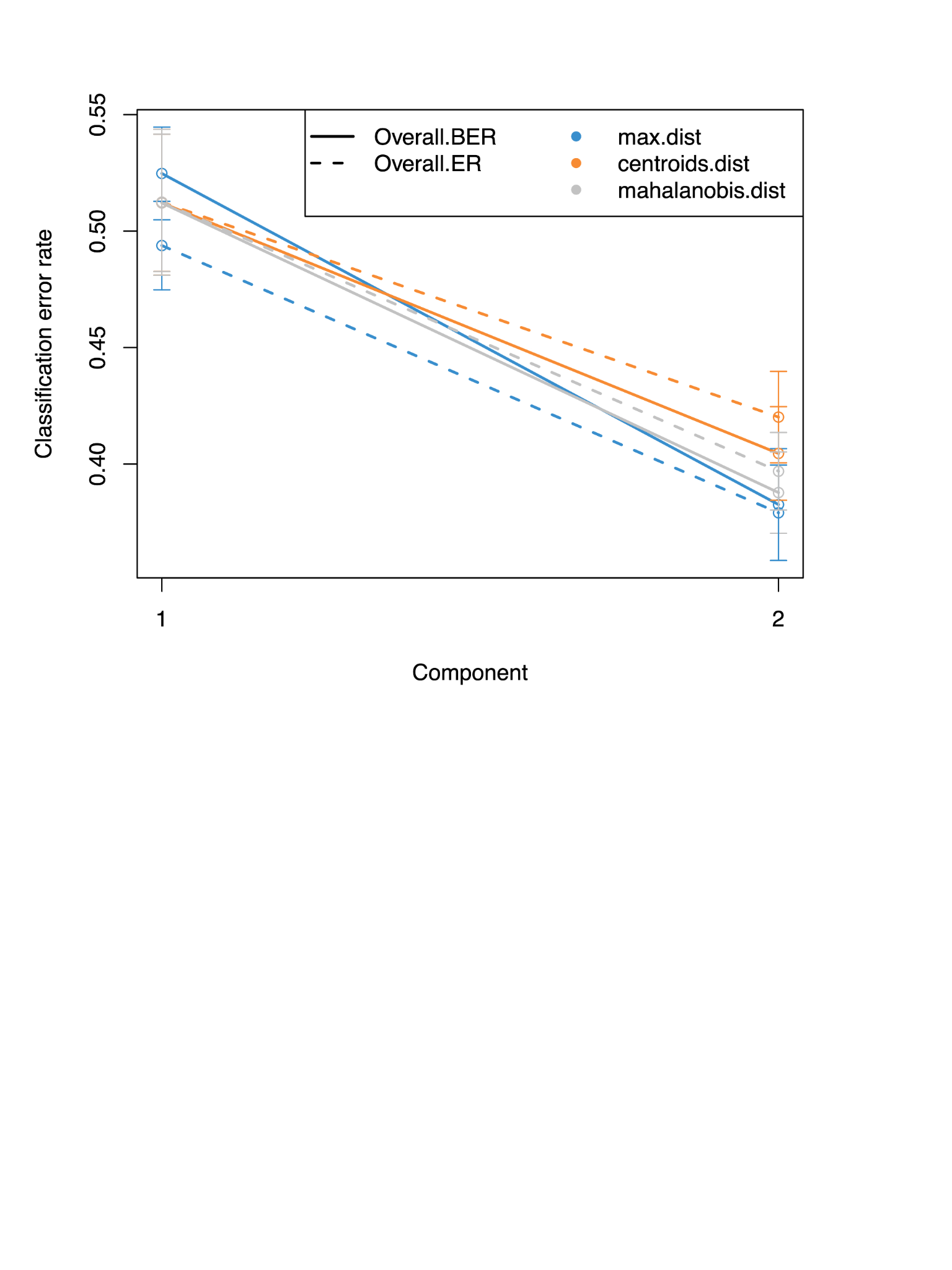

**Fig. S12 | Overall, per-disease and per-block classification error rates.** Error rate of each single-omics block (flow cytometry, cytokines, Treg, Teff), shown per disease and overall, for comparison with the integrated model. Bars are the mean and error bars the standard deviation across repeats.

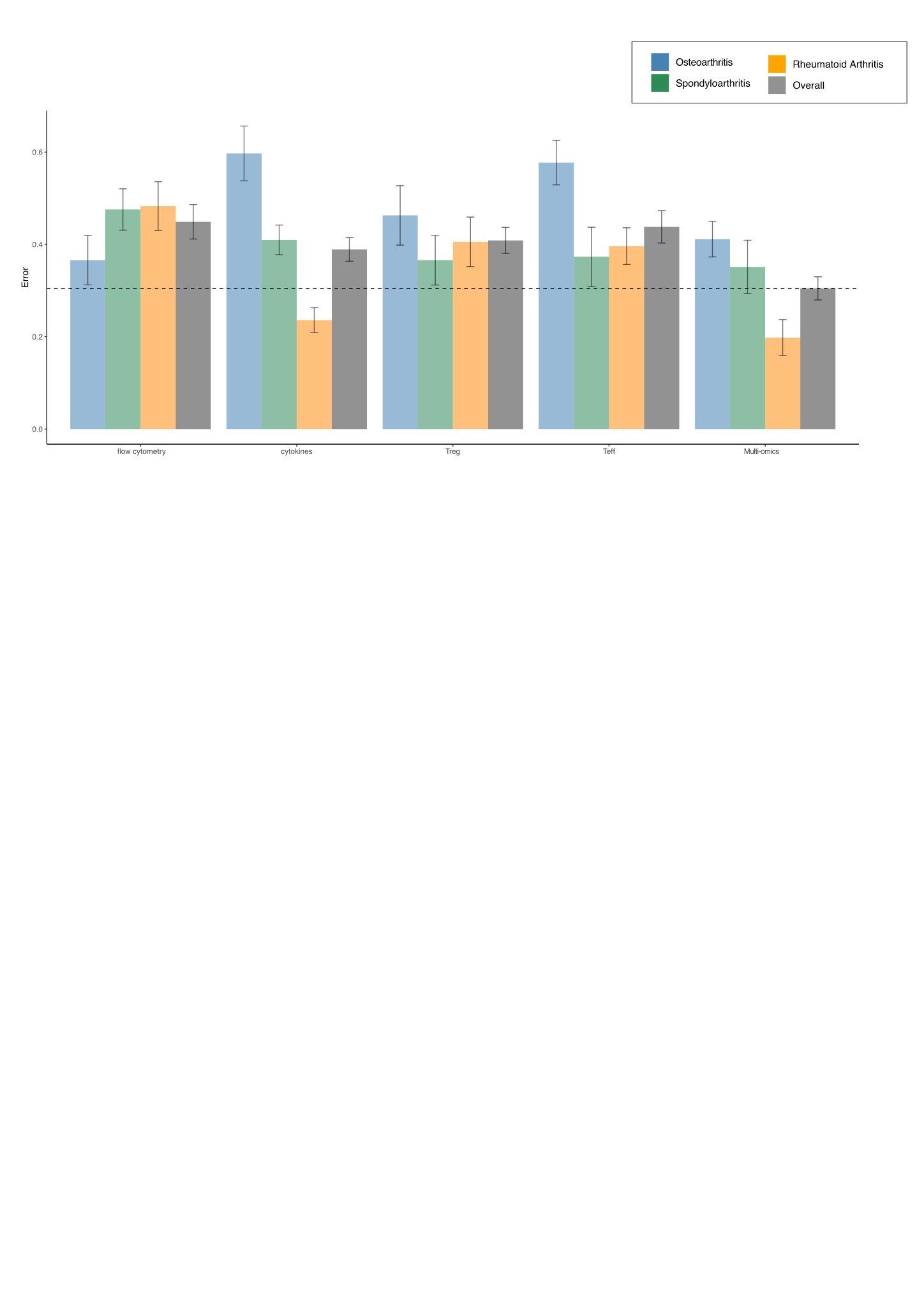

**Figure S13 | Per-block discrimination of each disease by ROC analysis.** One-versus-rest area under the ROC curve (AUC) for each omics block (flow cytometry, cytokines, Treg, Teff), computed on component 2 of the multi-block sPLS-DA model, for each disease class (OA vs rest, SpA vs rest, RA vs rest).

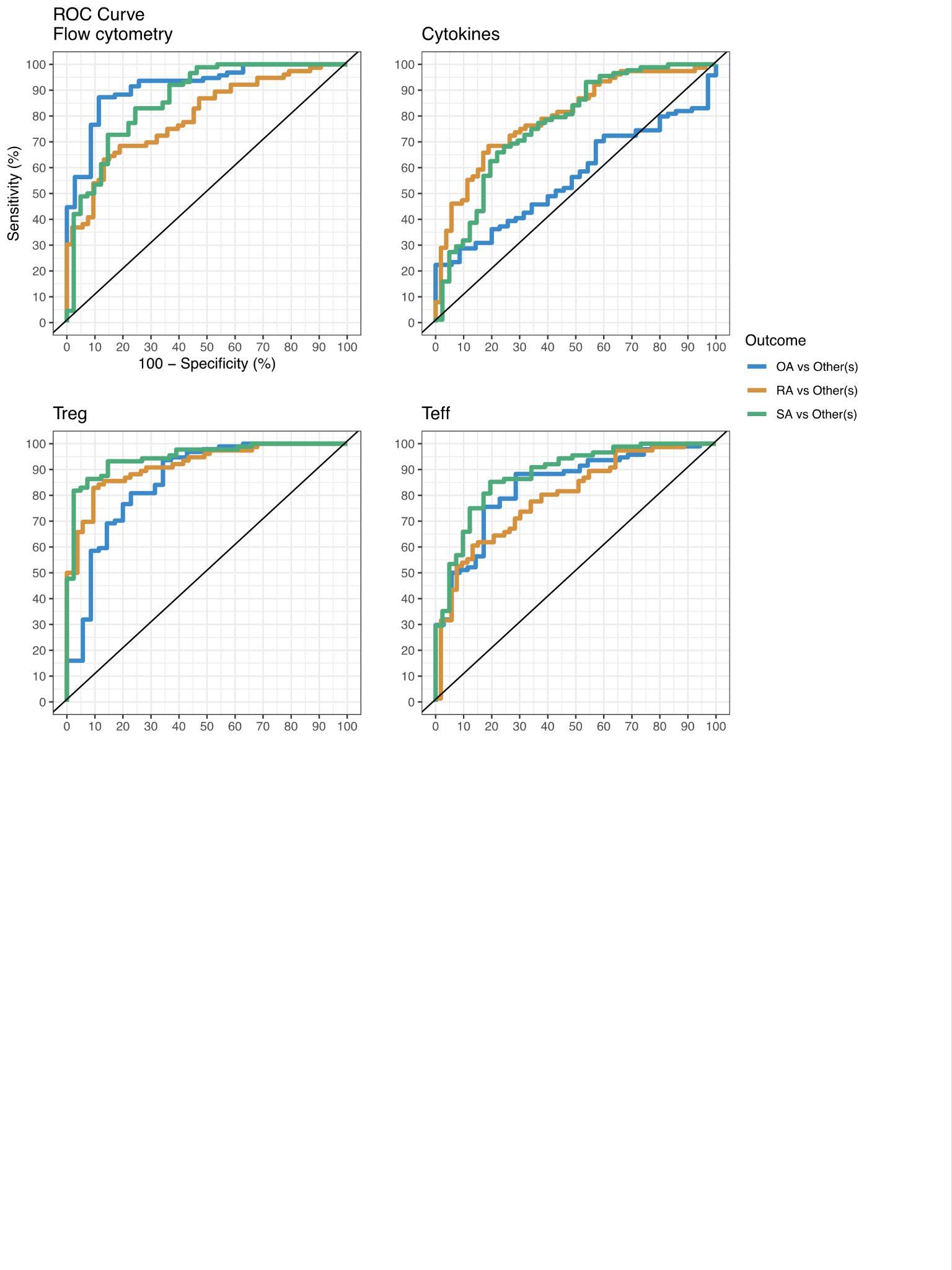

**Figure S14 | Side heatmap of the multi-omics analysis with clinical variables.**Clustered Image Map (CIM) heatmap of the multi-omics features selected on the first two components, displayed with clinical and demographic variables shown as side annotations: disease group, sex, age, BMI, erosive status, ACPA antibodies, rheumatoid factor (RF), DAS28-CRP, batch, early disease, HLA-B27, BASDAI, WOMAC, methotrexate, bDMARDs, and corticosteroids. Rows and columns were ordered by hierarchical clustering, with colour intensity reflecting the scaled value of each feature across samples.

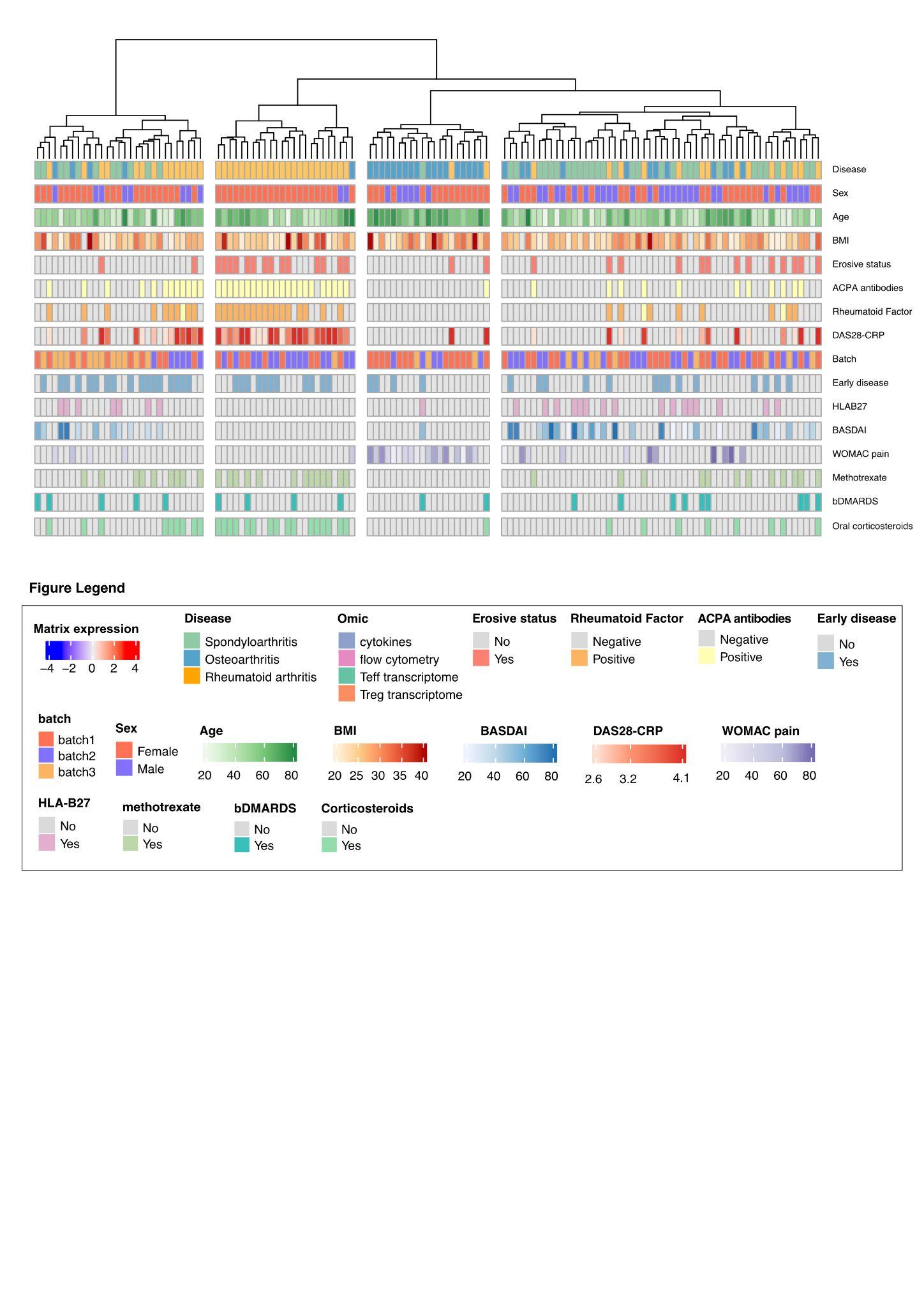

**Fig. S15 | Cluster composition by age and sex.** Distribution of age and sex across the four clusters in the whole cohort. Age is shown as the distribution of values per cluster (median and interquartile range), and sex as the proportion of female patients per cluster. Between-cluster differences were assessed using the Kruskal–Wallis test for age and Fisher's exact test for sex, with pairwise post hoc comparisons.

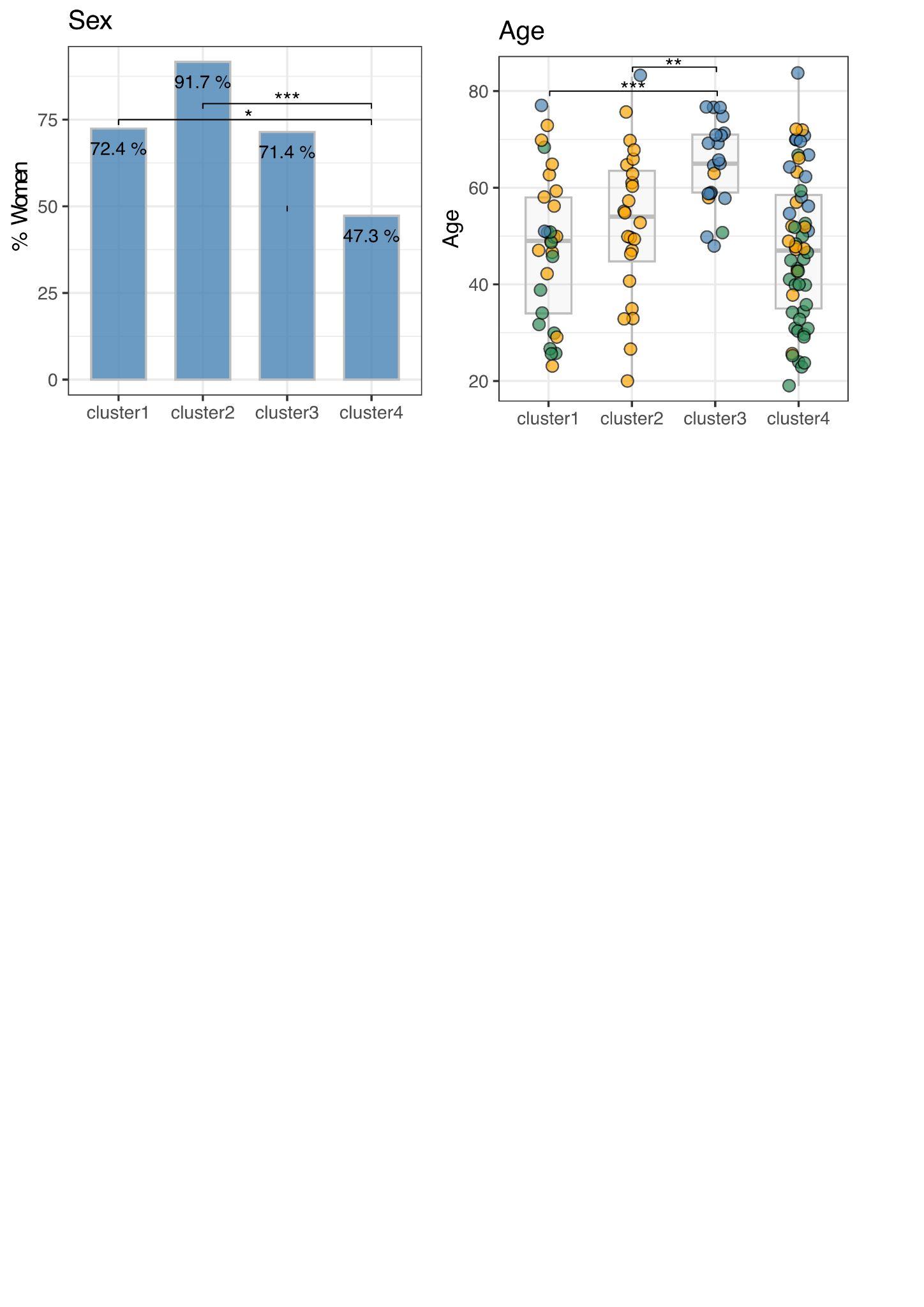

**Table S1 | Inclusion and Exclusion criteria of the Trans*i*mmunom study**

| **Inclusion criteria** | - **Age ≥ 18 years old** - **At least one auto-immune or auto-inflammatory disease including RA, SpAor knee OA (as a low grade inflammatory disease) according to ACR/EULAR classification criteria** |
| --- | --- |
|  | - **Additional criteria for OA:** - Uni or bilateral radiographic knee OA with a KL score of 2 or 3 (ACR criteria) - Patients ≥ 35 years old |
| **Exclusion criteria** | - **Pregnancy** - **Systemic infection in the month before the inclusion** - **Cancer and/or ongoing chemotherapy** - **Co existing infection by HBV HBC HIV** |
|  | - **Additional criteria for OA:** - KL score ≥ 4 - Co existing Inflammatory rheumatic disease (rheumatoid arthritis, spondyloarthritis, psoriasis) - Crystal induced arthropathies (chondrocalcinosis, gout) - Secondary OA (genetic disorder, hemochromatosis, avascular necrosis) - Contre-indication to X-rays |

**Table S2 | Rheumatoid arthritis clinical characteristic according to disease activity (N=91).** RA patients are stratified into remission–low disease activity (DAS28-CRP < 3.2; N = 37) and moderate–high disease activity (DAS28-CRP ≥ 3.2; N = 51). Continuous variables are expressed as mean and standard deviation (SD) within each subgroup, and categorical variables as percentages. Student's t-test and the chi-square test were applied, with significance defined as P ≤ 0.05. Missing values are expressed as percentages (%).

|  |  | **Rheumatoid Arthritis**  (N= 91) | **Remission - low disease activity**  (DAS28-CRP <3.2) (N=37) | **Moderate - high disease activity** (DAS28-CRP ≥3.2) (N=51) | p-value | missing  (%) |
| --- | --- | --- | --- | --- | --- | --- |
| **Age** (years) (mean (SD)) | | **49.71 (13.44)** | 46.30 (13.02) | 52.04 (13.45) | 0.048 | 0 |
| **BMI** (kg/m^2^) (mean (SD)) | | **25.79 (5.32)** | 25.18 (4.55) | 26.47 (5.77) | ns | 3.3 |
| **Sex** (%) | Female | **74 (81.3)** | 32 ( 86.5) | 39 (76.5) | ns | 0 |
|  | Male | **17 (18.7)** | 5 ( 13.5) | 12 (23.5) |  |  |
| **DAS28-CRP** (mean (SD)) | | **3.42 (1.28)** | 2.19 (0.58) | 4.31 (0.83) | <0.001 | 3.3 |
| **Erosions** (%) | | **47 (51.6)** | 19 ( 51.4) | 27 (52.9) | 0.672 | 1.1 |
| **Pulmonary RA** (%) | | **5 ( 5.5)** | 0 ( 0.0) | 5 ( 9.8) | 0.04 | 3.3 |
| **RF** (%) |  | **62 (68.1)** | 23 ( 62.2) | 37 (72.5) | ns | 4.4 |
| **ACPA** (%) |  | **81 (89.0)** | 35 ( 94.6) | 43 (84.3) | ns | 0 |
| **Oral corticosteroids** (%) | | **50 (54.9)** | 15 ( 40.5) | 34 (66.7) | 0.027 | 0 |
| **cDMARDs** (%) | | **62 (68.1)** | 27 ( 73.0) | 33 (64.7) | ns | 0 |
|  | Leflunomide | **4 ( 4.4)** | 2 ( 5.4) | 2 ( 3.9) |  |  |
|  | Methotrexate | **57 (62.6)** | 25 ( 67.6) | 30 (58.8) |  |  |
|  | Sulfasalazine | **1 ( 1.1)** | 0 ( 0.0) | 1 ( 2.0) |  |  |
| **bDMARDs** (%) | | **23 (25.3)** | 9 ( 24.3) | 13 (25.5) | ns | 0 |
|  | anti-TNF | **16 (17.6)** | 7 ( 18.9) | 8 (15.7) |  |  |
|  | anti-IL6 | **4 ( 4.4)** | 2 ( 5.4) | 2 ( 3.9) |  |  |
|  | Abatacept | **2 ( 2.2)** | 1 ( 2.7) | 1 ( 2.0) |  |  |
|  | JAK inhibitors | **1 ( 1.1)** | 0 ( 0.0) | 1 ( 2.0) |  |  |
|  | Rituximab | **2 ( 2.2)** | 0 ( 0.0) | 1 ( 2.0) |  |  |

**Table S3 | Spondyloarthritis clinical characteristic according to disease activity (N=58)** Spondyloarthritis patients are stratified by low disease activity (BASDAI < 40/100) (N=23) and high disease activity (BASDAI ≥ 40/100) (N=35). Continuous variables were expressed as mean and standard deviation (SD) in each subgroup, while categorical variables were expressed as percentages. A Student's t-test and chi-square test were applied and a significant p-value was defined with a threshold of ≤ 0.05. Missing values are expressed as a percentage (%).

|  |  | **Spondyloarthritis** (N=58) | **Low disease activity** (BASDAI < 40/100) (N=23) | **High disease activity** (BASDAI < 40/100) (N=35) | p-value | missing (%) |
| --- | --- | --- | --- | --- | --- | --- |
| **Age** (years) (mean (SD)) | | **39.50 (12.08)** | 40.43 (12.02) | 38.89 (12.26) | ns | 0 |
| **BMI** (kg/m^2^) (mean (SD)) | | **24.97 (4.50)** | 25.25 (4.56) | 24.79 (4.52) | ns | 3.4 |
| **Sex** (%) | Female | **27 ( 46.6)** | 10 ( 43.5) | 17 ( 48.6) | ns | 0 |
|  | Male | **31 ( 53.4)** | 13 ( 56.5) | 18 ( 51.4) |  |  |
| **Axial SpA**(%) | | **54 ( 93.1)** | 20 ( 87.0) | 34 ( 97.1) | ns | 0 |
| **Peripheral SpA**(%) | | **21 ( 36.2)** | 9 ( 39.1) | 12 ( 34.3) | ns | 3.4 |
| **Enthesitis SpA**(%) | | **35 (60.3)** | 10 (43.5) | 25 (71.4) | 0.044 | 3.4 |
| **HLA-B27** (%) | | **33 ( 56.9)** | 14 ( 60.9) | 19 ( 54.3) | ns | 15.5 |
| **Ankylosing Spondylitis** (%) | | **33 ( 56.9)** | 12 ( 52.2) | 21 ( 60.0) | ns | 12.1 |
| **Sacroiliitis MRI** (%) | | **39 ( 67.2)** | 15 ( 65.2) | 24 ( 68.6) | ns | 12.1 |
| **IBD** (%) | | **6 ( 10.3)** | 2 ( 8.7) | 4 ( 11.4) | ns | 0 |
|  | Crohn’s disease | **2 ( 3.4)** | 1 ( 4.3) | 1 ( 2.9) |  |  |
|  | Ulcerative colitis | **4 ( 6.9)** | 1 ( 4.3) | 3 ( 8.6) | ns | 0 |
| **Uveitis** (%) | | **8 ( 13.8)** | 4 ( 17.4) | 4 ( 11.4) | ns | 0 |
| **Reactive arthritis** (%) | | **3 ( 5.2)** | 0 ( 0.0) | 3 ( 8.6) | ns | 10.3 |
| **Dactylitis** (%) | | **8 ( 13.8)** | 2 ( 8.7) | 6 ( 17.1) |  | 6.6 |
| **Psoriasis** (%) | | **6 ( 10.3)** | 2 ( 8.7) | 4 ( 11.4) | ns | 8.7 |
| **BASDAI** (mean (SD)) | | **45.75 (21.86)** | 24.07 (12.72) | 60.00 (13.02) | <0.001 | 0 |
| **BASFI** (mean (SD)) | | **29.66 (25.95)** | 15.03 (20.83) | 39.28 (24.67) | <0.001 | 0 |
| **Oral NSAIDs** (%) | | **35 ( 60.3)** | 17 ( 73.9) | 18 ( 51.4) | ns | 8.6 |
| **cDMARDs** (%) | | **7 ( 12.1)** | 4 ( 17.4) | 3 ( 8.6) | ns | 0 |
|  | Methotrexate | **5 ( 8.6)** | 4 ( 17.4) | 1 ( 2.9) |  |  |
|  | Azathioprine | **1 ( 1.7)** | 0 ( 0.0) | 1 ( 2.9) |  |  |
|  | Sulfasalazine | **1 ( 1.7)** | 0 ( 0.0) | 1 ( 2.9) |  |  |
| **bDMARDs** (%) | | **6 ( 10.3)** | 3 ( 13.0) | 3 ( 8.6) | ns | 1.7 |
|  | Anti-TNF | **5 ( 8.6)** | 3 ( 13.0) | 2 ( 5.7) |  |  |
|  | Anti IL-12/IL-23 | **1 ( 1.7)** | 0 ( 0.0) | 1 ( 2.9) |  |  |

**Table S4 | Deep immunophenotyping cells populations and panels.** Cell populations were measured across 13 panels, including a general panel, B cells panel, NK-NCR cells panel, T cells activation and T cells naive/memory panel, MAIT-NKT cells panel, T cells migration panel, CD4 T cells polarization panel, T cells activation panel, Treg function panel, DC/monocytes panel, Treg phenotypes panel, and Myeloid-ILCs panel. The measurements were expressed in percentage (%) and include 137 cell populations across these panels.

| **Panel** | **Panel name** | **Variable name** | **Cell type** |
| --- | --- | --- | --- |
| Panel 01 | B cells | B cells (%) | B cells |
| Panel 01 | DC/monocytes | Monocytes (%) | Monocytes |
| Panel 01 | DC/monocytes | Classical Monocytes (%) | Monocytes |
| Panel 01 | DC/monocytes | Transient Monocytes (%) | Monocytes |
| Panel 01 | DC/monocytes | Resident Monocytes (%) | Monocytes |
| Panel 01 | Lymphoid | Lymphocytes (%) | Lymphocytes |
| Panel 01 | Memory T cells | Naive CD4 T cells (%) | CD4 T cells |
| Panel 01 | Myeloid - ILCs | Neutrophils/Eosinophils (%) | Neutrophils |
| Panel 01 | Myeloid - ILCs | Eosinophils (%) | Eosinophils |
| Panel 01 | Myeloid - ILCs | Neutrophils (%) | Neutrophils |
| Panel 01 | NK - NCR cells | NK cells (%) | NK cells |
| Panel 01 | NK - NCR cells | CD56dim CD16+ NK cells (%) | NK cells |
| Panel 01 | NK - NCR cells | CD56bright NK cells (%) | NK cells |
| Panel 01 | NK - NCR cells | CD56bright CD16+ NK cells (%) | NK cells |
| Panel 01 | T cells | T cells (%) | T cells |
| Panel 01 | T cells | NKT-like cells (%) | NKT cells |
| Panel 01 | T cells | CD4 T cells (%) | CD4 T cells |
| Panel 01 | T cells | CD8 T cells (%) | CD8 T cells |
| Panel 01 | T cells naive/memory | Central memory CD4 T cells (%) | CD4 T cells |
| Panel 01 | T cells naive/memory | Effector memory CD4 T cells (%) | CD4 T cells |
| Panel 01 | T cells naive/memory | TEMRA CD4+ cells (%) | CD4 T cells |
| Panel 01 | T cells naive/memory | Naive CD8 T cells (%) | CD8 T cells |
| Panel 01 | T cells naive/memory | Central memory CD8 T cells (%) | CD8 T cells |
| Panel 01 | T cells naive/memory | Effector memory CD8 T cells (%) | CD8 T cells |
| Panel 01 | T cells naive/memory | TEMRA CD8 T cells (%) | CD8 T cells |
| Panel 01 | Treg phenotypes | Treg CD25+ CD127+ cells (%) | CD4 T cells |
| Panel 02 | B cells | Naïve B cells (%) | B cells |
| Panel 02 | B cells | Unswitched memory B cells (%) | B cells |
| Panel 02 | B cells | Switched memory B cells (%) | B cells |
| Panel 02 | B cells | IgM+ IgD switched B cells (%) | B cells |
| Panel 02 | B cells | Plasmablast (%) | B cells |
| Panel 02 | B cells | Transitional B cells (%) | B cells |
| Panel 02 | B cells | CD21low B cells (%) | B cells |
| Panel 02 | B cells | CD10+ B cells (%) | B cells |
| Panel 02 | B cells | CD5+ B cells (%) | B cells |
| Panel 02 | B cells | Activated B cells CD32- (%) | B cells |
| Panel 03 | NK - NCR cells | CD56dim NKp44+ cells (%) | NK cells |
| Panel 03 | NK - NCR cells | CD56bright NKp44+ cells (%) | NK cells |
| Panel 03 | NK - NCR cells | CD56dim NKp30+ cells (%) | NK cells |
| Panel 03 | NK - NCR cells | CD56bright NKp30+ cells (%) | NK cells |
| Panel 03 | NK - NCR cells | CD56dim NKp46+ cells (%) | NK cells |
| Panel 03 | NK - NCR cells | CD56brightNKp46+ cells (%) | NK cells |
| Panel 03 | NK - NCR cells | CD56dim NKG2D+ cells (%) | NK cells |
| Panel 03 | NK - NCR cells | CD56bright NKG2D+ cells (%) | NK cells |
| Panel 03 | NK - NCR cells | CD56dim CD8+ NK cells (%) | NK cells |
| Panel 03 | NK - NCR cells | CD56bright CD8+ NK cells (%) | NK cells |
| Panel 03 | NK - NCR cells | CD56dim HLA-DR+ NK cells (%) | NK cells |
| Panel 03 | NK - NCR cells | CD56bright HLA-DR+ NK cells (%) | NK cells |
| Panel 03 | NK - NCR cells | NKG2D+ CD8 T cells (%) | CD8 T cells |
| Panel 03 | NK - NCR cells | HLA-DR+ CD8 T cells (%) | CD8 T cells |
| Panel 03 | NK - NCR cells | TCRgd T cells (%) | CD8 T cells |
| Panel 03 | NK - NCR cells | TCRgd CD16+ T cells (%) | TCRgd T cells |
| Panel 03 | NK - NCR cells | TCRgd NKG2D+ T cells (%) | TCRgd T cells |
| Panel 03 | NK - NCR cells | TCRgd HLA-DR+ T cells (%) | TCRgd T cells |
| Panel 03 | NK - NCR cells | TCRgd CD8+ T cells (%) | TCRgd T cells |
| Panel 03 | NK - NCR cells | NKT-like CD8+ (%) | NKT cells |
| Panel 03 | NK - NCR cells | NKT-like CD16+ cells (%) | NKT cells |
| Panel 03 | NK - NCR cells | NKT-like HLA-DR+ cells (%) | NKT cells |
| Panel 03 | NK - NCR cells | NKT-like NKG2D+ cells (%) | NKT cells |
| Panel 04 | T cells activation | CD8 T cells CD95+ (%) | CD8 T cells |
| Panel 04 | T cells activation | T effector cells CD95+ (%) | CD4 T cells |
| Panel 04 | T cells activation | Treg CD95+ (%) | CD4 T cells |
| Panel 04 | T cells activation | CD4 T cells CD95+ (%) | CD4 T cells |
| Panel 04 | T cells activation | CD8 T cells HLA-DR+ (%) | CD8 T cells |
| Panel 04 | T cells activation | T effector cells HLA-DR+ (%) | CD4 T cells |
| Panel 04 | T cells activation | Treg HLA-DR+ (%) | CD4 T cells |
| Panel 04 | T cells activation | CD4 T cells HLA-DR+ (%) | CD4 T cells |
| Panel 04 | T cells activation | CD8 ICOS+ (%) | CD8 T cells |
| Panel 04 | T cells activation | T effector cells ICOS+ (%) | CD4 T cells |
| Panel 04 | T cells activation | Treg ICOS+ (%) | CD4 T cells |
| Panel 04 | T cells activation | CD4 T cells ICOS+ (%) | CD4 T cells |
| Panel 04 | T cells naive/memory | CD4 T effector (%) | CD4 T cells |
| Panel 04 | T cells naive/memory | Naive T effector cells (%) | CD4 T cells |
| Panel 04 | T cells naive/memory | Central effector memory T cells (%) | CD4 T cells |
| Panel 04 | T cells naive/memory | Effector memory T cells (%) | CD4 T cells |
| Panel 04 | T cells naive/memory | TEMRA cells (%) | CD4 T cells |
| Panel 04 | T cells naive/memory | Naive T reg (%) | CD4 T cells |
| Panel 04 | T cells naive/memory | Central memory Treg (%) | CD4 T cells |
| Panel 04 | T cells naive/memory | Effector memory Treg (%) | CD4 T cells |
| Panel 04 | T cells naive/memory | TEMRA Treg (%) | CD4 T cells |
| Panel 05 | MAIT - NKT cells | NKT type 1 cells (%) | NKT cells |
| Panel 05 | MAIT - NKT cells | NKT TCR-Va7.2+ cells (%) | NKT cells |
| Panel 05 | MAIT - NKT cells | MAIT cells (%) | MCD4 T cells |
| Panel 07 | T cells migration | CD4 T cells CLA+ (%) | CD8 T cells |
| Panel 07 | T cells migration | CD8 T cells CLA+ (%) | CD4 T cells |
| Panel 07 | T cells migration | CD4 T cells CD49a+ (%) | CD8 T cells |
| Panel 07 | T cells migration | CD8 T cells CD49a+ (%) | CD4 T cells |
| Panel 07 | T cells migration | CD4 T cells CD103+ (%) | CD8 T cells |
| Panel 07 | T cells migration | CD8 T cells CD103+ (%) | CD4 T cells |
| Panel 07 | T cells migration | CD4 T cells a4+B7+ (%) | CD8 T cells |
| Panel 07 | T cells migration | CD8 T cells a4+B7+ (%) | CD4 T cells |
| Panel 07 | T cells migration | CD4 T cells a4+B7- (%) | CD8 T cells |
| Panel 07 | T cells migration | CD8 T cells a4+B7- (%) | CD8 T cells |
| Panel 08 | CD4 T cells polarization | TEMRA Th2 cells (%) | CD4 T cells |
| Panel 08 | CD4 T cells polarization | TEMRA Th17 cells (%) | CD4 T cells |
| Panel 08 | CD4 T cells polarization | TEMRA Th22 cells (%) | CD4 T cells |
| Panel 08 | CD4 T cells polarization | T helper Th2 cells (%) | CD4 T cells |
| Panel 08 | CD4 T cells polarization | T helper Th17 cells (%) | CD4 T cells |
| Panel 08 | CD4 T cells polarization | T helper Th1 cells (%) | CD4 T cells |
| Panel 08 | CD4 T cells polarization | T helper Th9-Th17.1 cells (%) | CD4 T cells |
| Panel 08 | CD4 T cells polarization | T helper Th22 cells (%) | CD4 T cells |
| Panel 08 | CD4 T cells polarization | Th1 memory cells (%) | CD4 T cells |
| Panel 08 | CD4 T cells polarization | Th2 memory cells (%) | CD4 T cells |
| Panel 08 | CD4 T cells polarization | Th17 memory cells (%) | CD4 T cells |
| Panel 08 | CD4 T cells polarization | Th22 memory cells (%) | CD4 T cells |
| Panel 08 | CD4 T cells polarization | TEMRA Th1 cells (%) | CD4 T cells |
| Panel 09 | T cells activation | CD4 T cells CD57+ (%) | CD4 T cells |
| Panel 09 | T cells activation | CD8 T cells CD57+ (%) | CD8 T cells |
| Panel 09 | T cells activation | CD4 T cells CD69+ (%) | CD4 T cells |
| Panel 09 | T cells activation | CD8 T cells CD69+ (%) | CD8 T cells |
| Panel 09 | T cells activation | CD4 T cells PD-1+ (%) | CD4 T cells |
| Panel 09 | T cells activation | CD8 T cells PD-1+ (%) | CD8 T cells |
| Panel 09 | T cells activation | CD4 T cells 4-1BB+ (%) | CD4 T cells |
| Panel 09 | T cells activation | CD8 T cells 4-1BB+ (%) | CD8 T cells |
| Panel 09 | T cells activation | CD4 T cells OX40+ (%) | CD4 T cells |
| Panel 09 | T cells activation | CD8 T cells OX40+ (%) | CD8 T cells |
| Panel 09 | T cells activation | CD4 T cells CD5hi (%) | CD4 T cells |
| Panel 09 | T cells activation | CD8 T cells CD5hi (%) | CD8 T cells |
| Panel 10 | Treg function | CD4 Treg CD39+ (%) | CD4 T cells |
| Panel 10 | Treg function | CD4 Treg GITR+ (%) | CD4 T cells |
| Panel 10 | Treg function | CD4 Treg LAP+ (%) | CD4 T cells |
| Panel 10 | Treg function | CD4 Treg LAG3+ (%) | CD4 T cells |
| Panel 10 | Treg function | CD4 Treg CD45RA- (%) | CD4 T cells |
| Panel 11 | DC/monocytes | pDCs (%) | DC/monocytes |
| Panel 11 | DC/monocytes | mDC1 (%) | DC/monocytes |
| Panel 11 | DC/monocytes | mDC2 (%) | DC/monocytes |
| Panel 11 | DC/monocytes | mDC CCR5+ (%) | DC/monocytes |
| Panel 11 | DC/monocytes | mDC (%) | DC/monocytes |
| Panel 11 | DC/monocytes | Basophils (%) | DC/monocytes |
| Panel 12 | Treg phenotypes | FoxP3+ Treg (%) | CD4 T cells |
| Panel 12 | Treg phenotypes | FoxP3+ Treg Helios+ (%) | CD4 T cells |
| Panel 12 | Treg phenotypes | FoxP3+ Treg CXCR5+ (%) | CD4 T cells |
| Panel 12 | Treg phenotypes | FoxP3+ Treg CTLA4+ (%) | CD4 T cells |
| Panel 13 | Myeloid - ILCs | Innate lymphoid cells (%) | ILCs |
| Panel 13 | Myeloid - ILCs | Innate lymphoid cells ILC2 (%) | ILCs |
| Panel 13 | Myeloid - ILCs | Innate lymphoid cells ILC1 (%) | ILCs |
| Panel 13 | Myeloid - ILCs | Innate lymphoid cells ILC3 (%) | ILCs |

CCR Chemokine receptor; CD, Cluster of differentiation; CLA, Cutaneous leukocyte-associated antigen; CTLA-4, Cytotoxic T-lymphocyte–associated antigen; DC dendritic cells, FoxP3, Forkhead box P3; GITR, Glucocorticoid-Induced TNFR-Related; HLA-DR, Human leukocyte antigen-DR isotype; ICOS, Inducible costimulator; ILC, Innate lymphoid cells; Ig, Immunoglobulin; LAG-3, Lymphocyte activation gene 3; LAP, LC3-associated phagocytosis, MAIT, Mucosal-associated invariant T; NK, Natural killer; NKG2D, Natural killer group 2 member D; PD-1, Programmed Cell Death Protein; TCR, T-cell receptor, TEMRA, Terminally differentiated effector memory cell; Treg, T regulatory cell

**Table S5 | Cytokines and Panel measured in the Trans*i*mmunom study.** Cytokines were measured using four Luminex panels including Cytokines-Chemokines kit (CC-CK), Highly sensitive kit (HS), Receptor kit, and TH17 kit. Measurements were expressed in MFI (mean fluorescence intensity) and included 62 cytokines

| **Panel** | **Cytokines abbreviation** | **Cytokines** |
| --- | --- | --- |
| CCCK | EGF | Epidermal Growth Factor |
| CC-CK | Eotaxin | Eotaxin |
| CC-CK | FGF2 | Fibroblast Growth Factor 2 |
| CC-CK | Fractalkine | Fractalkine |
| CC-CK | GCSF | Granulocyte Colony-Stimulating Factor |
| CC-CK | GMCSF | Granulocyte-Macrophage Colony-Stimulating Factor |
| CC-CK | GRO | Growth-Regulated Oncogene |
| CC-CK | IFNa2 | Interferon Alpha 2 |
| CC-CK | IFNg | Interferon Gamma |
| CC-CK | IL10 | Interleukin 10 |
| CC-CK | IL12p40 | Interleukin 12p40 |
| CC-CK | IL12p70 | Interleukin 12p70 |
| CC-CK | IL13 | Interleukin 13 |
| CC-CK | IL15 | Interleukin 15 |
| CC-CK | IL1a | Interleukin 1 Alpha |
| CC-CK | IL1b | Interleukin 1 Beta |
| CC-CK | IL1RA | Interleukin 1 Receptor Antagonist |
| CC-CK | IL2 | Interleukin 2 |
| CC-CK | IL3 | Interleukin 3 |
| CC-CK | IL4 | Interleukin 4 |
| CC-CK | IL5 | Interleukin 5 |
| CC-CK | IL6 | Interleukin 6 |
| CC-CK | IL7 | Interleukin 7 |
| CC-CK | IL8 | Interleukin 8 |
| CC-CK | IL9 | Interleukin 9 |
| CC-CK | IP10 | Interferon-gamma-Inducible Protein 10 |
| CC-CK | MCP1 | Monocyte Chemoattractant Protein 1 |
| CC-CK | MCP3 | Monocyte Chemoattractant Protein 3 |
| CC-CK | MDC | Macrophage-Derived Chemokine |
| CC-CK | MIP1a | Macrophage Inflammatory Protein 1 Alpha |
| CC-CK | MIP1b | Macrophage Inflammatory Protein 1 Beta |
| CC-CK | sCD40L | Soluble CD40L |
| CC-CK | TGFa | Transforming Growth Factor Alpha |
| CC-CK | TNFa | Tumor Necrosis Factor Alpha |
| CC-CK | TNFb | Tumor Necrosis Factor Beta |
| CC-CK | VEGF | Vascular Endothelial Growth Factor |
| Receptor | sCD30 | Soluble CD30 |
| Receptor | sEGFR | Epidermal Growth Factor Receptor |
| Receptor | sgp130 | Soluble gp130 |
| Receptor | sIL1R1 | Interleukin 1 Receptor Type 1 |
| Receptor | sIL1R2 | Interleukin 1 Receptor Type 2 |
| Receptor | sIL2Ra | Interleukin 2 Receptor Alpha |
| Receptor | sIL4R | Interleukin 4 Receptor |
| Receptor | sIL6R | Interleukin 6 Receptor |
| Receptor | sRAGE | Receptor for Advanced Glycation End Products |
| Receptor | sTNFR1 | Tumor Necrosis Factor Receptor Type 1 |
| Receptor | sTNFR2 | Tumor Necrosis Factor Receptor Type 2 |
| Receptor | sVEGFR1 | Vascular Endothelial Growth Factor Receptor 1 |
| Receptor | sVEGFR2 | Vascular Endothelial Growth Factor Receptor 2 |
| Receptor | sVEGFR3 | Vascular Endothelial Growth Factor Receptor 3 |
| TGFBeta | TGFB1 | Transforming Growth Factor Beta 1 |
| TH17 | IL17A | Interleukin 17A |
| TH17 | IL17E | Interleukin 17E |
| TH17 | IL17F | Interleukin 17F |
| TH17 | IL21 | Interleukin 21 |
| TH17 | IL22 | Interleukin 22 |
| TH17 | IL23 | Interleukin 23 |
| TH17 | IL27 | Interleukin 27 |
| TH17 | IL28A | Interleukin 28A |
| TH17 | IL31 | Interleukin 31 |
| TH17 | IL33 | Interleukin 33 |
| TH17 | MIP3a | Macrophage Inflammatory Protein 3 alpha |

CC-CK, Cytokines - Chemokines kit; HS Highly sensitive; TH, T helper

**Table S6 | Cell populations associated with rheumatologic diseases (N = 36).** Analyses were performed on the Transimmunom cohort (N = 240), comprising HV (N = 47), OA (N = 44),RA patients (N = 91), and SpA) patients (N = 58). Differences between each disease group and the Healthy Volunteer reference group were assessed using the non-parametric Wilcoxon rank-sum test (Mann-Whitney U test). P values were adjusted for multiple testing using the Benjamini-Hochberg (BH) procedure to control the false discovery rate. Only cell populations showing significant differences (p ≤0.05) are shown.

**Table S7 | Cytokines associated with rheumatologic diseases (n=47).** Analyses were performed on 239 Transimmunom participants (HV, N = 47; OA, N = 44; RA, N = 91; Sp A, N = 57). Each disease group was compared with HV (reference group) using the Mann–Whitney U test. Only cytokines showing significant differences (p ≤ 0.05) are shown.

**Table S8 | Differentially expressed genes in Teffs across disease groups versus HV (n = 177).** Complete list of differentially expressed genes identified in CD4+ effector T cells (Teff) for each pairwise comparison (RA (N=82) vs HV (N=38), SpA (N=51) vs HV, and OA (N=44) vs HV). Genes were considered differentially expressed at an absolute log2 fold change exceeding log2(2) and a Benjamini-Hochberg FDR-adjusted p-value below 0.05. Columns indicate gene symbol, base mean normalised count, log2 fold change (MLE), log2 fold change standard error, Wald statistic, nominal p-value, adjusted p-value, and disease comparison.

**Table S9 | Differentially expressed genes in Treg across disease groups versus HV (n=262).** Complete list of differentially expressed genes identified in CD4+ regulatory T cells (Treg) for each pairwise comparison (RA (N=59) vs HV (N=22), SpA (N=45) vs HV, and OA (N=35) vs HV). Genes were considered differentially expressed at an absolute log2 fold change exceeding log2(2) and a Benjamini-Hochberg FDR-adjusted p-value below 0.05. Columns indicate gene symbol, base mean normalised count, log2 fold change (MLE), log2 fold change standard error, Wald statistic, nominal p-value, adjusted p-value, and disease comparison.

**Table S10 | Over-representation analysis of Gene Ontology Biological Process terms in Teff and Treg cells across disease groups.** Results of over-representation analysis (ORA) performed using clusterProfiler against Gene Ontology Biological Process (GO-BP) terms, for CD4+ effector T cells (Teff) and CD4+ regulatory T cells (Treg) independently. For each disease group versus healthy volunteers comparison (RA vs HV, SpA vs HV, OA vs HV), nominally significant genes (raw p-value ≤ 0.05) were stratified by direction of effect and tested separately for upregulated (log2FC > 0) and downregulated (log2FC < 0) gene sets against a background universe comprising all genes detected and retained after low-count filtering in the respective DESeq2 model. Enrichment was assessed with Benjamini-Hochberg multiple testing correction and a q-value threshold of 0.2. Redundant GO terms were removed by semantic similarity-based simplification (similarity cutoff = 0.7, retention by minimum adjusted p-value). Columns indicate GO term identifier, GO term description, gene ratio, background ratio, nominal p-value, Benjamini-Hochberg adjusted p-value, q-value, gene count, gene symbols of contributing genes, cell type (Teff or Treg), direction of regulation (up or down), and disease comparison.

**Table S11 | Discriminatory performance (AUC) of immunological measurement modalities for rheumatic diseases.** Area under the receiver operating characteristic curve (AUC) values are shown for one-versus-rest classification of each arthritis subtype (OA, SpA, RA) against all other groups combined. Performance is reported separately for four measurement modalities: flow cytometry, cytokine profiling, regulatory T cells (Treg), and effector T cells (Teff).

|  | **Flow cytometry** | **Cytokines** | **Treg** | **Teff** |
| --- | --- | --- | --- | --- |
| OA vs Other(s) | 0.916 (p=4.459e-13) | 0.5635 (p=0.2683) | 0.8553 (p=5.938e-10) | 0.8377 (p=3.984e-09) |
| SpA vs Other(s) | 0.864 (p=3.005e-11) | 0.7697 (p=8.587e-07) | 0.9484 (p=2.22e-16) | 0.8814 (p=3.404e-12) |
| RA vs Other(s) | 0.797(p=1.061e-08) | 0.7984 (p=8.709e-09) | 0.9163 (p=8.882e-16) | 0.7949 (p=1.292e-08) |

**Table S12 | Multi-omics feature loadings in component 1 discriminating osteoarthritis, spondyloarthritis, and rheumatoid arthritis.**

Feature-level results from an integrative multi-omics analysis across three disease groups. Each row is a measured feature; the importance column gives its loading (weight) on the latent component, and the OA/SpA/RA columns give the mean value of that feature within each disease group. The GroupContrib column assigns each feature to the disease group in which its mean is maximal. Features are ordered by descending absolute importance within each modality.

**Table S13 | Multi-omics feature loadings in component 2 discriminating osteoarthritis, spondyloarthritis, and rheumatoid arthritis.**

Feature-level results from an integrative multi-omics analysis across three disease groups. Each row is a measured feature; the importance column gives its loading (weight) on the latent component, and the OA/SpA/RA columns give the mean value of that feature within each disease group. The GroupContrib column assigns each feature to the disease group in which its mean is maximal. Features are ordered by descending absolute importance within each modality.

**Table S14 | Composition of the four clusters according to clinical and biological characteristics, overall and by disease group.** Patients were grouped into four clusters and are presented for the whole cohort (Global, N = 129) and separately for each diagnosis: (RA, N = 53) (SpA, N = 41), (OA, N = 35). Continuous variables are expressed as median (IQR) and categorical variables as percentage (N). Continuous variables were compared across clusters using the Kruskal–Wallis test, and categorical variables using Fisher's exact test. *p* < 0.05 was considered statistically significant; ns = not significant.

| **Global (N=129)** | **cluster 1 (N=29)** | **cluster 2**  **(N=24)** | **cluster 3**  **(N=21)** | **cluster 4 (N=55)** | **p -value** | **NA (%)** |
| --- | --- | --- | --- | --- | --- | --- |
| Age (years) | 49 (34-58) | 54 (44.75-63.5) | 65 (59-71) | 47 (35-58.5) | p <0.0001 | 0 |
| Sex (%) | 72.4% (N=21/29) | 91.7% (N=22/24) | 71.4% (N=15/21) | 47.3% (N=26/55) | p=0.0007 | 3.9 |
| BMI (kg/m2) | 26.1 (22.4-30.1) | 25.0 (23.0-29.9) | 28.1 (21.2-32.5) | 25.9 (22.1-28.4) | ns | 0 |
| hs CRP (mg/L) | 4.74 | 3.8 (2.2-12.3) | 2.1 (0.7-3.6) | 2.5 (0.7-6.6) | ns | 15.5 |
| early disease (%) | 58.6 % (N=17/29) | 45.8 % (N=11/24) | 20% (N=4/20) | 26.4% (N=14/53) | p = 0.009 | 2.3 |
| **RA (N=53)** | **cluster(N=14)** | **cluster 2 (N=23)** | **cluster 3 (N=2)** | **cluster 4 (N=14)** | **p -value** |  |
| Erosive status (%) | 14.3% (N=2/14) | 63.6% (N=14/22) |  | 92.9% (N=13/14) | p <0.0001 |  |
| ACPA (%) | 92.3% (N=12/13) | 95.6% (N=22/23) |  | 78.6% (N=11/14) | ns | 1.9 |
| RF (%) | 76.9% (N=10/13) | 69.6% (N=16/23) |  | 57.1% (N=8/14) | ns | 1.9 |
| DAS 28 - CRP (0-9.8) | 3.5 (2.8-4.1) | 3.8 (3.0-4.2) |  | 3.2 (2.4-4.5) | ns | 1.9 |
| oral corticosteroids (%) | 71.4% (N=10/14) | 69.6% (N=16/22) |  | 57.1% (N=8/14) | ns | 0 |
| Methotrexate (%) | 64.2% (N=9) | 56.5 (N=13) |  | 85.7% (N=12) | ns | 0 |
| bDMARDS (%) | 28.6% (N=4) | 17.4% (N=4) |  | 35.7% (N=5) | ns | 0 |
| **SpA (N=41)** | **Cluster (N=11)** | **cluster 2 (N=0)** | **cluster 3 (N=1)** | **cluster 4 (N=29)** | **p -value** |  |
| bDMARDS (%) | 9.1% (N=1/11) |  |  | 14.8% (N=4/27) | ns | 4.9 |
| BASDAI (0-10) | 43.5 (37.5-56) |  |  | 48.9 (31.4-66) | ns | 2.4 |
| HLA-B27 (%) | 87.5% (N=7/8) |  |  | 69.6% (N=16/23) | ns | 21.9 |
| **OA (N=35)** | **cluster (N=4)** | **cluster 2 (N=1)** | **cluster 3 (N=18)** | **cluster 4 (N=12)** | **p -value** |  |
| WOMAC pain (0-100) |  |  | 45 (40-65) | 65 (40-71.3) | ns | 2.9 |

**Table S15 | Post-hoc pairwise comparisons between clusters for variables showing a significant overall difference.** For each variable that reached statistical significance in the overall between-cluster analysis (Table S14), pairwise comparisons were performed between each pair of clusters. Continuous variables (Age) were compared using pairwise Wilcoxon rank-sum (Mann–Whitney) tests, and categorical variables (Sex, Early disease, Erosive status) using Fisher's exact test. p-values are reported for each cluster pair; ns = not significant (p ≥ 0.05).

| **Variable** | **group1** | **group2** | **p-value** |
| --- | --- | --- | --- |
| Age | cluster2 | cluster1 | ns |
| Age | cluster3 | cluster1 | 0.0003 |
| Age | cluster4 | cluster1 | ns |
| Age | cluster3 | cluster2 | 0.0091 |
| Age | cluster4 | cluster2 | ns |
| Age | cluster4 | cluster3 | 0.0001 |
| Sex | cluster1 | cluster2 | ns |
| Sex | cluster1 | cluster3 | ns |
| Sex | cluster1 | cluster4 | 0.037 |
| Sex | cluster2 | cluster3 | ns |
| Sex | cluster2 | cluster4 | 0.0001 |
| Sex | cluster3 | cluster4 | ns |
| Early disease | cluster1 | cluster2 | ns |
| Early disease | cluster1 | cluster3 | 0.009 |
| Early disease | cluster1 | cluster4 | 0.008 |
| Early disease | cluster2 | cluster3 | ns |
| Early disease | cluster2 | cluster4 | ns |
| Early disease | cluster3 | cluster4 | ns |
| Erosive status | cluster1 | cluster2 | 0.0057 |
| Erosive status | cluster1 | cluster3 | 0.05 |
| Erosive status | cluster1 | cluster4 | <0.0001 |
| Erosive status | cluster2 | cluster3 | ns |
| Erosive status | cluster2 | cluster4 | ns |
| Erosive status | cluster3 | cluster4 | ns |

**Table S16 | Disease duration across clusters: additional comparison supporting the early-disease finding.** As early disease emerged as a key discriminating feature between clusters (Table S14), disease duration was examined as a complementary continuous measure across the four clusters (Global cohort, N = 129). Values are expressed as median (IQR) in years. Although the overall Kruskal–Wallis test across all four clusters was not significant, cluster 1, characterized by a higher proportion of early disease, was taken as the reference and compared pairwise against the other clusters using Wilcoxon rank-sum (Mann–Whitney) tests. Disease duration was significantly shorter in cluster 1 than in cluster 3 (p = 0.001) and showed a trend toward being shorter than in cluster 4 (p = 0.058), whereas the difference with cluster 2 was not significant. This pattern is consistent with, and reinforces, the categorical early-disease result reported in Table S14.

| **Global (N=129)** | **cluster 1 (N=29)** | **cluster 2**  **(N=24)** | **cluster 3**  **(N=21)** | **cluster 4 (N=55)** | **Wilcoxon p -value** |
| --- | --- | --- | --- | --- | --- |
| Disease duration (years) | 2.0 (1.0 - 5.0) | 5.0 (0.8 - 9.2)  ns | 5.0 (3.0 - 7.2)  (p=0.001) | 5.0 (2.0-10.0) | p =0.058 |

**Table S17 | Markov Cluster (MCL) algorithm partitioning of the early-disease signature protein–protein interaction network.**

Proteins encoded by the differentially expressed genes were mapped onto a STRING protein–protein interaction network and partitioned into functional modules using the Markov Cluster (MCL) algorithm. For each resulting cluster, the number of member genes, the primary functional description of the module, and the constituent protein names are shown. Cluster 1 (14 proteins) was enriched for interleukin-10 signalling; cluster 4 (2 proteins) corresponded to the Regulator of G protein signalling (RGS) domain family. Clusters 2 and 3 had no significantly enriched primary annotation ("–").

| **cluster number** | **gene count** | **primary description** | **protein names** |
| --- | --- | --- | --- |
| 1 | 14 | Interleukin-10 signaling | CD24, TNF, SLC11A1, IL1RN, PTGS2, CXCL2, IER3, TNFRSF12A, BCL2A1, PTX3, ADRB2, PNP, ABCA1, CLDN5 |
| 2 | 9 | - | GZMH, MS4A1, BLNK, CA2, TCL1A, SPIB, IRF8, MZB1, FCRLA |
| 3 | 2 | - | GLI1, CHKA |
| 4 | 2 | Regulator of G protein signalling domain | RGS18, RGS16 |

**Table S18 | Functional enrichment analysis of the early-disease signature.**

Over-representation analysis of the early-disease signature genes was performed using STRING against the DISEASES, KEGG and Reactome annotation databases. For each enriched term, the table reports the category, term identifier and description, the number of observed versus background genes, the enrichment strength (log₁₀ of observed/expected), the signal score, the false discovery rate (Benjamini–Hochberg corrected), and the matching proteins in the network (Ensembl protein IDs and gene labels). Only terms passing a false discovery rate < 0.05 are shown. Enriched terms included immune system disease, primary immunodeficiency disease and autoimmune disease (DISEASES), NF-κB signalling (KEGG), and interleukin-10 signalling (Reactome).
